# Molecular basis of UHRF1 allosteric activation for synergistic histone modification binding by PI5P

**DOI:** 10.1101/2021.08.04.455045

**Authors:** Papita Mandal, Zhadyra Yerkesh, Vladlena Kharchenko, Levani Zandarashvili, Dalila Bensaddek, Lukasz Jaremko, Ben E. Black, Wolfgang Fischle

## Abstract

Chromatin marks are recognized by distinct binding modules many of which are embedded in multidomain proteins or complexes. How the different protein functionalities of complex chromatin modulators are regulated is often unclear. Using a combination of biochemical, biophysical and structural approaches we delineated the regulation of the H3unmodified and H3K9me binding activities of the multidomain UHRF1 protein. The phosphoinositide PI5P interacts with two distant flexible linker regions of UHRF1 in a mode that is dependent on the polar head group and the acyl part of the phospholipid. The associated conformational rearrangements stably position the H3unmodified and H3K9me3 histone recognition modules of UHRF1 for multivalent and synergistic binding of the H3 tail. Our work highlights a novel molecular function for PI5P outside of the context of lipid mono- or bilayers and establishes a molecular paradigm for the allosteric regulation of complex, multidomain chromatin modulators by small cellular molecules.

## Main

Conformational flexibility and its control are a hallmark of biological regulation. Especially, intrinsically disordered regions allow proteins to explore a wide range of conformational space enabling interaction with different partners and function in diverse contexts^1,2^. Deterministic changes in protein conformation often occur in a regulated manner. Triggers for protein conformational change include differential splicing, posttranslational modification and allosteric ligand binding. While the general importance of these regulatory modes has long been recognized, the exact molecular details of their working mechanisms are often unclear.

The specific recognition of chromatin modifications, i.e. DNA methylation and histone marks is a hallmark of epigenetic regulatory processes. While a large number of protein domains and folds have now been described to work in this context, their function and regulation in multidomain proteins and multiprotein complexes is still poorly understood.

We have previously reported that the multidomain chromatin effector and writer protein UHRF1 (Ubiquitin-like with PHD and RING finger domains 1) is an allosteric target of the phosphoinositide PI5P that modulates the chromatin engagement of different histone binding domains of the factor^3^. UHRF1 acts as a safeguard of the genome by maintaining global DNA methylation levels, protecting chromatin from DNA damaging agents, maintaining higher-order chromatin structures and silencing repetitive DNA elements^4-10^. Accordingly, the protein is up-regulated in various types of malignancies^11-14^.

UHRF1 is composed of five domains that are connected through flexible linkers (Fig. 1a). The ubiquitin-like domain (UBL) directs ubiquitylation activity of UHRF1 towards histone H3^15,16^. A tandem tudor domain (TTD) and a plant homeodomain (PHD) recognize methylation at lysine 9 (K9me) and the unmodified N-terminus of the histone H3 tail, respectively^17-19^. The SET- and RING-associated (SRA) domain preferentially binds to hemimethylated (CpG) DNA^20,21^ and the Really Interesting New Gene (RING) domain catalyzes H3 ubiquitination on K18 and/or K23^22-24^. While the functions of the individual domains of UHRF1 are well defined, their interplay is only emerging. Here, communication between the interdomain linker regions seems to be crucial^3,25-28.^ For example, hemimethylated DNA triggers a not yet molecularly defined activation of UHRF1 that results in cooperation of the UBL and RING domains for enhanced H3 ubiquitylase function^15,16,24,29-33^. Further, we have demonstrated that a polybasic region (PBR) in the Linker 4 between SRA and RING domains can block TTD-H3K9me binding. Allosteric binding of PI5P releases the PBR from the TTD enabling its H3K9me recognition^3^. Other work unveiled that differential splicing in the flexible linker regions affects the overall regulation of human and mouse UHRF1^26^. The complex regulation of UHRF1 has made this factor an attractive model to study how a multidomain chromatin protein is conformationally and functionally controlled.

**Fig. 1:**
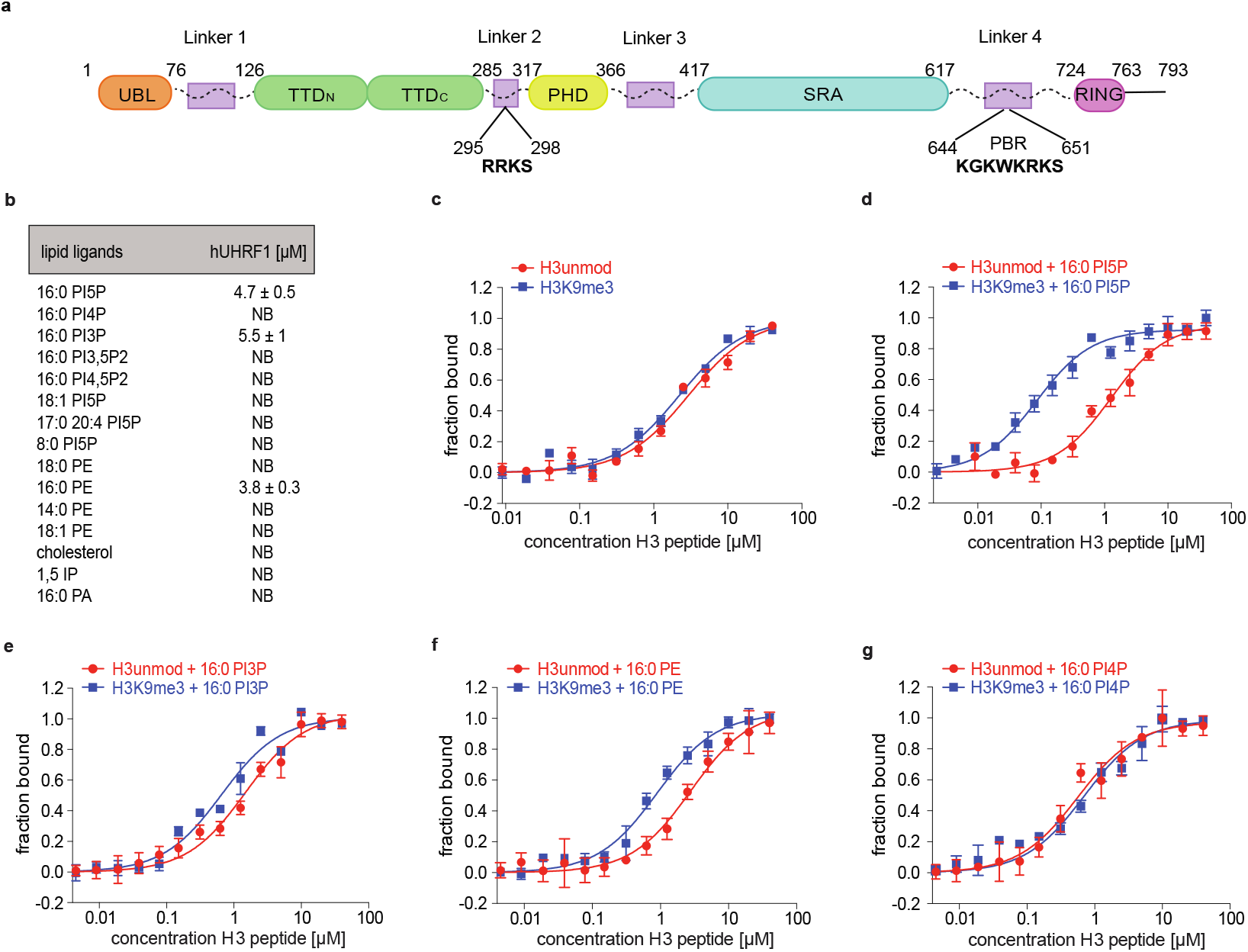
hUHRF1 recognizes PIPs with a monophosphorylated inositol headgroup and di-C 16:0 acyl chains but gets activated only by PI5P. **a**, Scheme illustrating the domain architecture of hUHRF1. UBL, ubiquitin-like domain; TTD, tandem tudor domain (TTDN-TTDC); PHD, plant homeodomain; SRA, SET and RING associated domain; RING, really interesting new gene domain. Four putatively flexible linker regions connect the structured domains. PBR, polybasic region. Patches of amino acids relevant for this study are annotated. **b**, Binding of hUHRF1 to phospholipids and other lipids of the indicated composition was analyzed by microscale thermophoresis. *K*D deduced as average from three independent titration measurements is listed. Error corresponds to SD. NB, not binding. **c** to **g**, Titration series of H3 peptides (residues 1-20) of the indicated modification state with fluorescently labeled recombinant hUHRF1 were analyzed by microscale thermophoresis in apo-state (c) and in presence of different phospholipids (d-g). Data are plotted as average of three independent experiments; error bars correspond to SD. K_D_ values of all measurements are listed in Extended Data Table 1.

Here, we set out to delineate the molecular details of PI5P-mediated allosteric activation of human UHRF1 (hUHRF1). While hUHRF1 discriminates for the configuration of the phosphoinositide acyl chains, it binds different mono-phosphorylated inositol head groups (i.e., PI3P, PI5P). However, our biochemical and structural data show that only PI5P is functional in establishing a composite binding mode that bridges two inter-domain linkers of hUHRF1. The resulting conformational rearrangement establishes multivalent, synergistic recognition of the H3K9me tail by the TTD-PHD module.

## Results

### hUHRF1 recognizes the inositol head group and acyl chains of phosphoinositides

To better understand the molecular details of PI5P binding, we first determined the specificity of hUHRF1 for different phospholipids. In quantitative, in-solution measurements using microscale thermophoresis (MST), recombinant, full-length hUHRF1 (Extended Data Fig. 1a) displayed a strong preference for mono-vs. bis (3,5 or 4,5)-phosphorylated inositol head groups with only phosphoinositides carrying phosphorylation in the C3 (K_D_ (PI3P) = 5.5 μM) and C5 (K_D_ (PI5P) = 4.7 μM) positions showing interaction. Notably, the low micromolar binding to PI5P is right in the range of the predicted and indirectly measured nuclear concentration of this phosphoinositide^34,35^. Remarkably, only PI5P carrying the di-C16:0 but not the shorter di-C 8:0 or longer unsaturated 17:0,20:4 or di-C 18:1 acyl chain configuration bound hUHRF1 in these assays (Fig. 1b). Neither inositol 1,5-bisphosphate (1,5-IP) representing the isolated head group of PI5P nor di-C 16:0 phosphatidic acid (16:0 PA) representing the isolated tail-group of PI5P were bound by hUHRF1. Of the different tested phospholipids only the di-C 16:0 form of phosphatidylethanolamine (16:0 PE) but not its di-C 14:0, di-C 18:0 nor the di-C 18:1 counterparts showed interaction. Also, the unrelated lipid cholesterol was not bound by hUHRF1. Overall, the results indicated that hUHRF1 interacts with and discriminates the whole phosphoinositide molecule with specificity for the phosphorylation of the inositol head group and selectivity for the lengths and saturation level of the acyl chains.

### PI5P specifically enhances hUHRF1 H3K9me binding

Next, we analyzed the effect of phospholipids on the interaction of hUHRF1 with the H3 tail. We focused exclusively on the di-C 16:0 forms, as only these had shown significant interaction with the protein. As Fig. 1c,d shows, PI5P only mildly (∼ 2.6 fold) increased the binding strength of hUHRF1 to an unmodified H3 tail peptide (K_D_(apo-hUhRF1) = 2.9 μM, K_D_(hUHRF1(PI5P)) = 1.2 μM). In contrast, interaction with a corresponding H3K9me3 peptide was increased up to 27-fold with the PI5P bound hUHRF1 displaying one of the strongest affinities to this modification observed (K_D_ = 0.08 μM) (see Extended Data Tables 1 and 2 for a full listing of K_D_s determined in this study). On the contrary, PI3P and PE, which showed interaction with hUHRF1 comparable to PI5P, only induced a mild discriminatory effect on H3 tail recognition (Fig. 1e,f). PI4P that did not interact with hUHRF1 in the direct binding studies did not have any effect on the binding to the H3 tail (Fig. 1g).

Previous reports have shown hemimethylated DNA and the UBL1-2 domain of USP7 influence H3 tail binding of hUHRF1^27,33,36^. Compared to PI5P we detected only mild enhancement (3-4 fold difference) of hUHRF1 interaction with H3K9me3 in the presence of both these ligands (Extended Data Fig. 1b,c and Extended Data Table 1). Based on these analyses we concluded that PI5P (di-C 16:0) is a unique and most potent allosteric activator of hUHRF1 H3K9me3-binding.

### PI5P directs multivalent, synergistic interaction of hUHRF1 with H3K9me3

To understand the differential peptide-binding modes of apo- and PI5P-bound hUHRF1, we performed hydrogen-deuterium exchange (HDX) mass spectrometry (MS) using recombinant hUHRF1. This method has been successfully employed to demonstrate allosteric regulation of multidomain proteins by measuring the changes in backbone structure and dynamics (e.g., PARP1)^37,38^. We compared the HDX pattern of the different forms of hUHRF1 in the absence and presence of unmodified and H3K9me3 peptides. HDXMS experiments were performed over a time course of 10^1^ to 10^4^ s and revealed a number of partially overlapping peptides that were protected from deuterium uptake upon H3 peptide binding. For all experiments, almost no change in deuterium uptake was observed in regions outside of the TTD and PHD (Fig. 2a and Extended Data Figs. 2, 3 and 4). In the apo-state the HDXMS patterns observed for hUHRF1 were highly similar for the unmodified and K9me3 H3 tails. The peptides with decreased deuterium exchange mapped in both cases primarily to a region of the PHD (region II, residues 313-357 aa) (Extended Data Figs. 2b and 3b) that according to 3D structural studies interfaces directly with the N-terminus of H3^18,19^. In contrast, peptides covering the TTD region showed similar deuterium exchange kinetics in the absence and in the presence of the unmodified and K9me3 H3-tail (Fig. 2a-e). The results were consistent with the TTD of hUHRF1 not contributing to H3-tail binding in the absence of PI5P. Interaction in this state is solely mediated by the PHD.

**Fig. 2:**
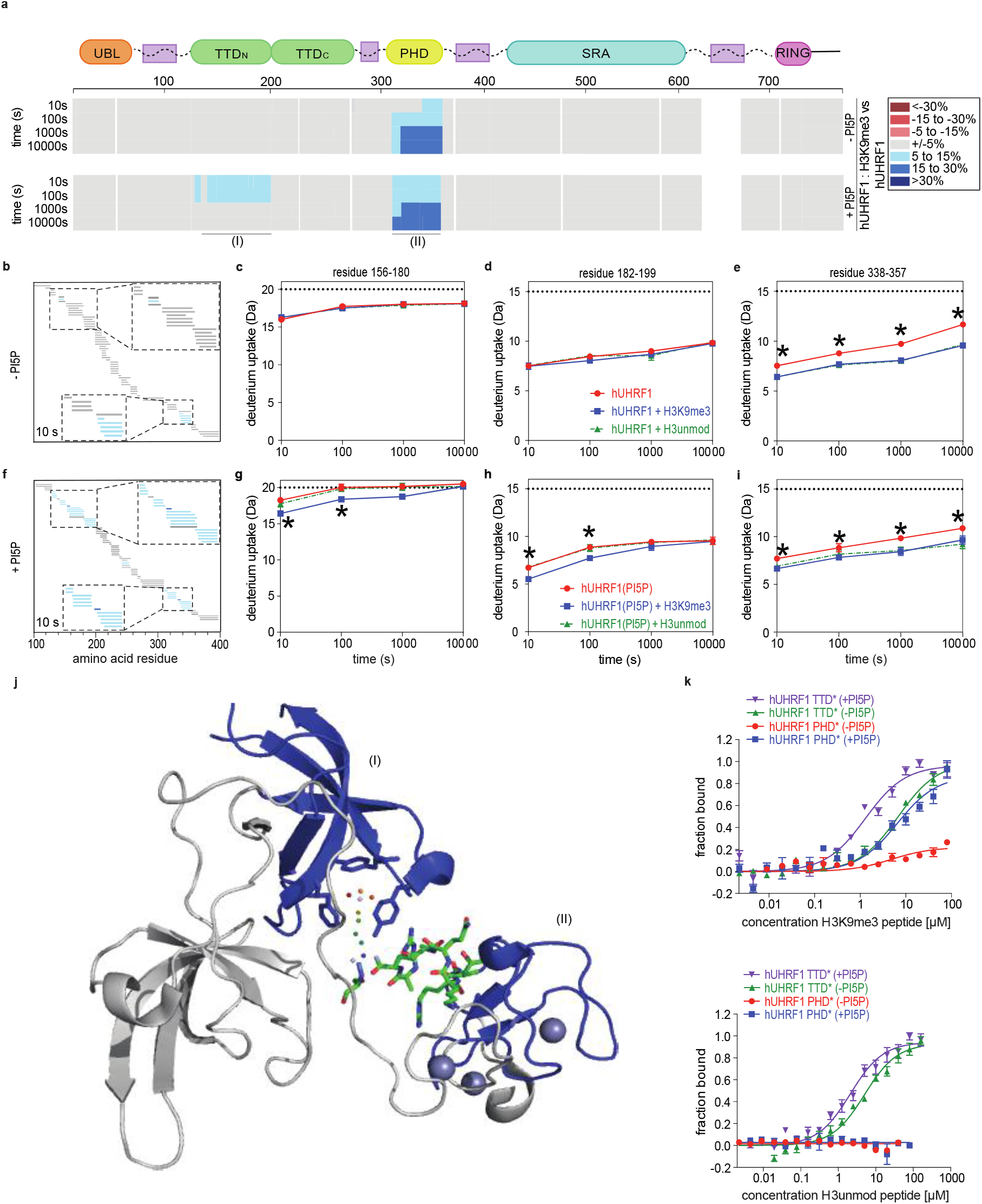
PI5P directs multivalent, synergistic interaction of the hUHRF1 TTD-PHD module with H3K9me3. **a**, The consensus at each amino acid position of the percentual difference of HDX at each time point is plotted with and without PI5P for each of >200 unique partially overlapping peptides of the hUHRF1/H3K9me3 complex (see Extended Data Fig. 2). The color key for binning of HDX differences is shown at the right. White regions indicate gaps in the obtained peptide coverage of hUHRF1. Roman numerals indicate regions highlighted in the PDB structure 4GY5 (see panel j). **b**, Percentual difference in HDX upon binding to H3K9me3 peptide at 10 s along the TTD-PHD region of hUHRF1. Each small horizontal bar represents a peptide in the dataset representing the indicated region of hUHRF1. Coloring indicates HX difference upon binding. Insets show enlargement of the marked regions. **c** to **e**, HDX time curves of specific peptides encompassing the aromatic cage residues of the TTD domain (c and d) and the N-terminal H3 tail-interacting residues in the PHD region (e) of apo-hUHRF1 binding the H3K9me3 peptide. Each experiment was performed in triplicate; asterisks indicate time points with P < 0.01 (Student’s t test between hUHRF1 and hUHRF1/H3K9me3 experiments). Dashed line reflects full deuteration level of the corresponding peptide. Error bars represent SD from three independent measurements. **f**, Same analysis as in (b) but in presence of PI5P. **g** to **i**, Same analysis as in (c-e) but in presence of PI5P. **j**, Crystal structure of the TTD-PHD module in complex with H3K9me3 peptide (PDB code 4GY5). Regions undergoing differential HDX are numbered and highlighted in blue. **k**, Titration series of H3K9me3 (upper panel) and H3 unmodified (lower panel) peptides with full-length hUHRF1 carrying mutations in the TTD or PHD domains in absence and presence of di-C 16:0 PI5P. TTD*: Y188A, Y191H; PHD*: D334, D337A. Data are plotted as average of three independent experiments; error bars correspond to SD. K_D_ values are listed in Extended Data Table 1.

In the PI5P-bound state, the unmodified H3-tail caused a HDXMS pattern similar to the apo-state of hUHRF1 with peptides in the PHD but not the TTD region being protected from deuterium exchange (Extended Data Fig. 3a). In contrast, the K9me3 H3-tail caused significant decrease in deuterium uptake in the PHD (region II) but also the TTD (region I) (Fig. 2a, f-j and Extended Data Fig. 2a). Importantly, peptides spanning the aromatic cage residues of the TTD (152-200 aa) were found to be protected. The findings implied that hUHRF1 recognizes the H3K9me3 mark specifically via the TTD domain in the PI5P bound state.

Different reader domains of composite proteins or complexes can either work individually/independently or in combination (multivalent and/ or synergistic) with each other^26,39,40^. The overall binding strength is higher in multivalent, synergistic interactions compared to independent multivalent binding (reflecting only the sum of the individual domain binding strength). To determine whether the enhanced binding of hUHRF1 to H3K9me3 in the presence of PI5P is due to independent (i.e. TTD and PHD working separately) or synergistic (i.e. TTD and PHD working in concert) multivalency, we measured H3 peptide-binding affinities of hUHRF1 after mutating either TTD or PHD and in absence and presence of PI5P. Of importance for these experiments, neither the previously described TTD (TTD*: Y188A/Y191H)^17^ nor PHD (PHD*: D334/D337A)^41^ mutations had any effect on hUHRF1 PI5P binding (Extended Data Fig. 5a and Extended Data Table 2). While hUHRF1 PHD* in the apo-state did not bind the unmodified or K9me3 H3-tail, interaction with the H3K9me3 peptide was observed in the PI5P-bound state (Fig. 2k). The results were in agreement with the TTD being blocked in the absence of PI5P and contributing to H3-tail binding in the presence of PI5P.

When analyzing hUHRF1 TTD*, this mutant recognized both unmodified (K_D_ = 5 μM) and K9me3 H3-tail peptides (K_D_ = 6.7 μM) with similar affinities, likely via the functional and free PHD domain. However, unlike the wild type protein this mutant showed only slight enhancement of interaction with the H3 unmodified (K_D_= 2.1 μM) and H3K9me3 (K_D_ = 1.3 μM) peptides in the presence of PI5P. We note that the binding strength for H3K9me3 was significantly higher (K_D_ = 80 nM) for PI5P-bound, wild type hUHRF1 (Fig. 1c and Extended Data Table 1) compared to the sum of the contribution of the individual TTD and PHD. Collectively, the findings pointed to PI5P binding establishing a TTD-PHD dependent multivalent, synergistic recognition mode of hUHRF1 (Fig.2j).

### PBR/ Linker 4 is necessary but not sufficient for PI5P-binding of hUHRF1

In previous work we showed that PI5P binds to the polybasic region (PBR) of hUHRF1. While this interaction unblocks the TTD from PBR-mediated inhibition^3^, it cannot explain the observed multivalent, synergistic binding of H3K9me3 by hUHRF1 in the presence of PI5P. Additional changes in the protein that functionally couple TTD and PHD must occur. To uncover the molecular mechanism behind PI5P driven enhanced H3K9me3 recognition, we first aimed to further characterize the PI5P binding of hUHRF1.

As shown previously^3^, deletion of the C-terminal region (617-793 aa) of hUHRF1 caused complete loss of interaction with PI5P. Using a series of deletion constructs (not shown), we mapped the phospholipid-binding region to Linker 4 (605-675 aa) that bound PI5P with a similar affinity as the intact protein (Fig. 3a). The binding specificity of this fragment was comparable to that of full-length hUHRF1 with strong preference for di-C 16:0 monophosphorylated phospholipids (Extended Data Fig. 5b).

**Fig. 3:**
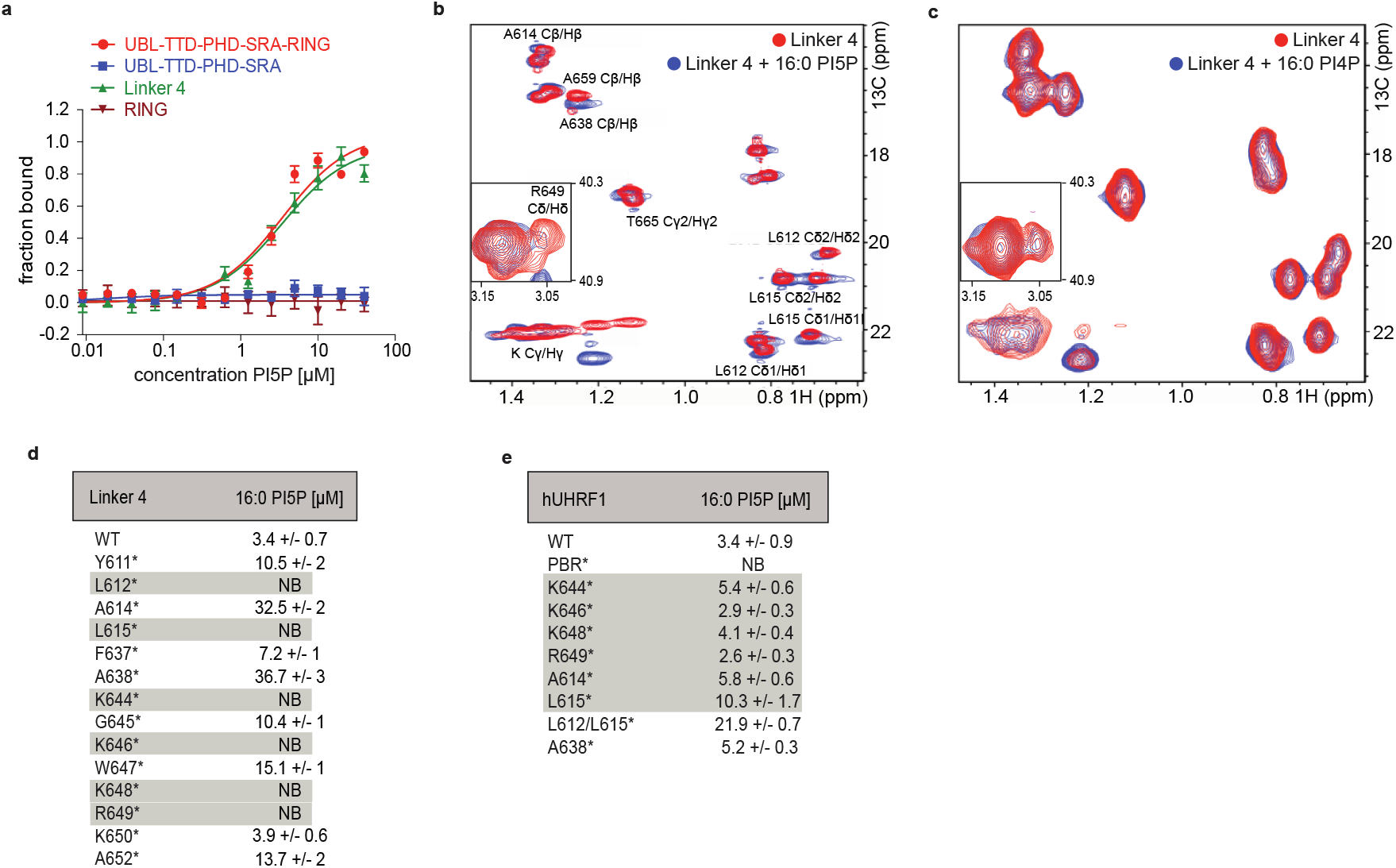
The PI5P binding mode of PBR/Linker 4 is different from full-length hUHRF1. **a**, Titration series of di-C 16:0 PI5P with fluorescently labeled, recombinant, full length and truncated hUHRF1 analyzed by microscale thermophoresis. Data are plotted as average of three independent experiments; error bars correspond to SD. K_D_ values are listed in Extended Data Table 2. **b** and **c**, Solution NMR spectra of hUHRF1 Linker 4 (residues 605-675) in absence and presence of di-C 16:0 PI5P (2D ^1^ H-^13^ C HSQC-CT (constant time)) (b) and di-C 16:0 PI4P 2D ^1^ H-^13^ C HSQC) (c). Amino acids that showed changes in chemical shift upon PI5P binding are indicated by residue number/type. The overlapped signals of the aliphatic ^1^ H/^13^ C correlations of lysine and arginine (enlarged in the inset) side chains are shown in (b). **d** and **e**, Dissociation constants (*K*D) for di-C 16:0 PI5P binding of wild type and point mutated recombinant Linker 4 (d) and full-length hUHRF1 (e) as determined by microscale thermophoresis. Mutations showing significant effects are highlighted. PBR*: K644A, K646A, K648A, R649A, K650A, S651A. Results represent averages of minimally three independent experiments. NB, not binding; error represents SD.

To obtain further insights into the interaction of PI5P with Linker 4, we employed high-resolution multidimensional solution NMR spectroscopy. Comparison of ^1^H-^13^C HSQC (Heteronuclear Single Quantum Coherence spectroscopy) spectra of Linker 4 in the absence and presence of di-C 16:0 PI5P showed significant chemical shift changes (Fig. 3b). To rule out that these chemical shift changes were unspecific and putatively caused by adding a phospholipid to the Linker 4 polypeptide, we repeated the measurements using di-C 16:0 PI4P. No differences in the chemical shifts compared to Linker 4 alone were observed (Fig. 3c). We managed to fully assign the Linker 4 resonance peaks using a combination of 3D triple resonance ^1^H-^15^ N-detected experiments (targeting backbone and side-chain resonances) and ^1^H-^13^C-detected 3D ^13^C-edited NOESY and (H)CCH TOCSY spectra (completing the side-chain assignments). The experiments identified two classes of Linker 4 residues being selectively affected by PI5P and not by PI4P binding (Fig. 3b). The first class consists of hydrophobic side-chains, i.e., A614, A659, A638, T665, L612 and L615, and the second class encompasses positively charged residues. While in contrast to the well-resolved hydrophobic side-chains containing methyl groups we could not resolve all of the positively charged residues due to severe overlap of the aliphatic ^1^H-^13^C correlations of lysine and arginine side chains, positively charged residues displayed considerable chemical shifts and signal shape changes indicative of the interaction (Fig. 3b, see the Cδ/Hδ region of R649 overlapped with other arginines (zoomed on a small panel) and Cγ/Hγ region of the methyl groups of lysines on the main spectrum).

Solution-state structure determination of Linker 4 in the PI5P bound state was not possible due to the lipid-protein complex aggregating at the concentrations required for this analysis. We thus resorted to analyzing point mutants of Linker 4 targeting the residues that showed chemical shift changes in the presence of the phosphoinositide. Besides mutation of the hydrophobic residues L612 and L615, mutation of the charged residues K644, K646, K648 and R649 but not K650 completely abolished interaction of Linker 4 with PI5P (Fig. 3d). Other residues like A614, A638 showed significant attenuation in interaction with the respective K_D_s reduced by ca. 10-fold. We then set out to test these mutations in the context of full-length hUHRF1. To our surprise, none of the individual mutations that abolished the binding of PI5P to Linker 4 had a similar effect in the context of the full-length protein. Mutation of the charged resides K644, K646, K648 and R649 had virtually no effect, while mutation of L615 and the double mutant L612/L615 showed some loss of interaction (3- and 6-fold lower K_D_ compared to wild type hUHRF1, respectively) (Fig. 3e and Extended Data Table 2). As only combination of mutation of multiple Linker 4 residues (K644, K646, K646 and R649) completely abolished the binding of PI5P (we refer to this mutant as PBR*), we postulated that additional binding interfaces must exist for this phospholipid in full-length hUHRF1.

### PI5P mediates interaction of hUHRF1 Linker 2 and Linker 4

Multiple attempts to obtain X-ray crystals of full-length hUHRF1 in absence and presence of PI5P failed. It also turned out impossible to analyze hUHRF1/PI5P binding in detail using NMR due to the high concentrations of protein and in particular phospholipid required for this method. We therefore turned to HDXMS which requires significantly less concentrated hUHRF1/PI5P complex to obtain structural insights into this interaction. In the initial analysis we compared the HDX pattern of PI5P-bound hUHRF1 with the protein in the apo-state. Even though we didn’t get full coverage for the PBR and surrounding regions, we noticed strong protection from deuterium uptake in peptides covering Linker 4 and a region immediately upstream corresponding to the C-terminus of the SRA domain (580-605 aa) (Extended Data Fig. 6a). However, a low level of protection was observed throughout the protein and we were unable to map any other regions affected by PI5P. We reasoned that the presence of a lipid moiety is putatively causing desolvation of the protein during HDX and thereby masking HDX patterns caused by PI5P binding. To map out such background effects, we took advantage of the interaction of hUHRF1 with di-C 16:0 PI3P, which while being specific and mediated by Linker 4 does not enhance hUHRF1 H3K9me3-binding (Fig. 1 and Extended Data Fig. 5b). Comparing the HDX patterns of the PI5P bound protein with those of the PI3P bound protein indeed allowed further mapping of the molecular consequences of hUHRF1 PI5P binding.

HDXMS experiments were performed over a time course of 10^1^ to 10^5^ s (Fig. 4a and Extended Data Fig. 7). As predicted, the Linker 4 and C-terminal region of the SRA did not show differential protection in the analysis, since both, PI5P and PI3P interact with these regions. We observed significant decrease in HDX in the TTD domain (127-151 aa; region I), in Linker 2 (274-312 aa; region II) and in the PHD domain (319-340 aa; region III) (Fig. 4a-d,f and Extended Data Fig. 7a). These regions either participate directly in PI5P binding and/ or undergo conformational rearrangements that minimize their deuterium uptake.

**Fig. 4:**
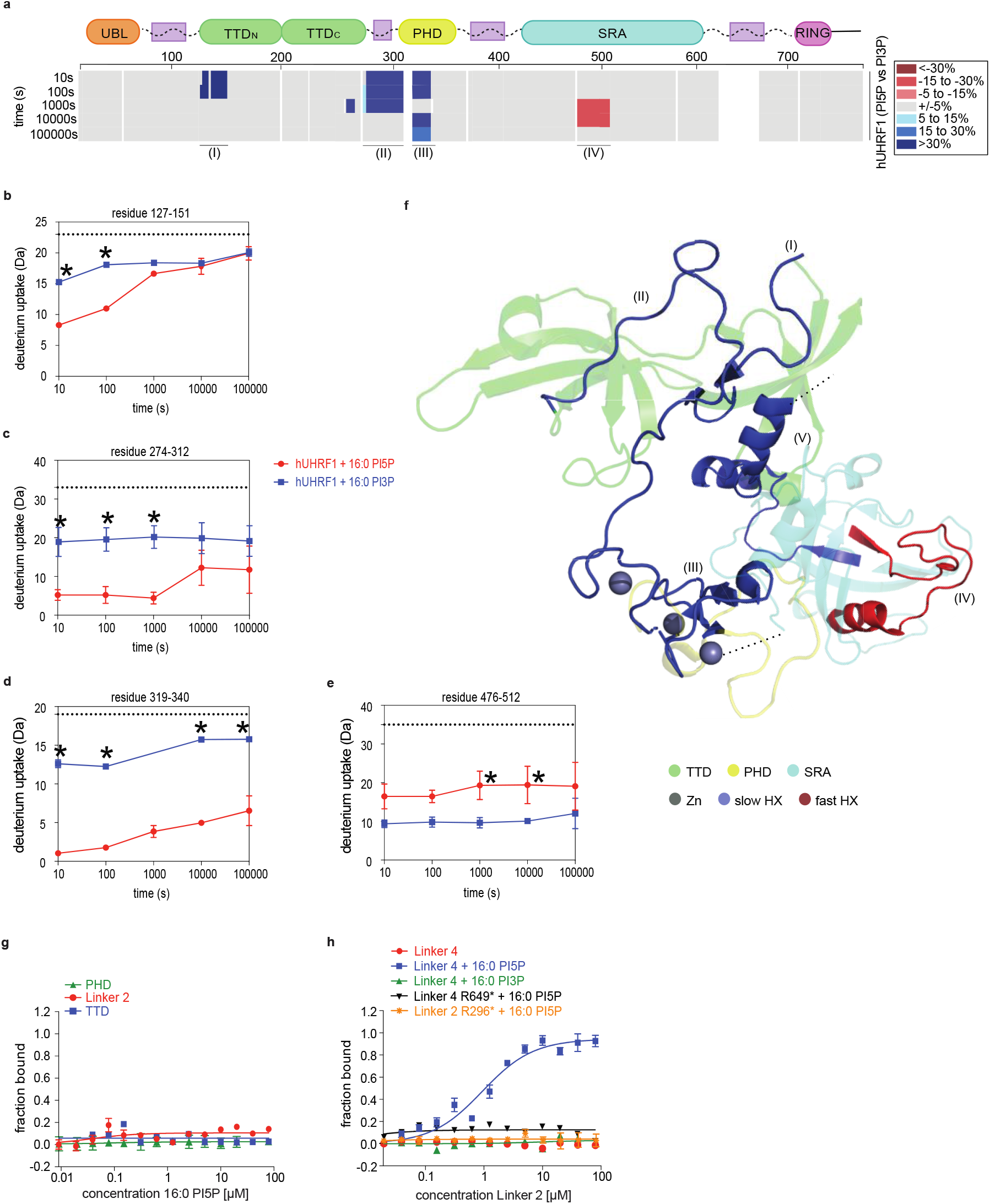
PI5P is a conformation-specific allosteric activator of hUHRF1. **a**, The consensus at each amino acid position of the percentual difference of HDX at each time point is plotted with and without PI5P for each of >200 unique partially overlapping peptides of hUHRF1. The color key for binning of HDX differences is shown at the right. White regions indicate gaps in the obtained peptide coverage of hUHRF1. Roman numerals indicate regions highlighted in the PDB structure in (f). Scheme illustrating domain architecture of hUHRF1 is at the top. **b** to **e**, HDX time curves of specific peptides encompassing the TTD surface groove (b), the Linker 2 (c), the N-terminal H3 tail-interacting residues of the PHD (d) and the DNA binding region of the SRA (e) are shown. Each experiment was performed in triplicate; asterisks indicate time points with P < 0.01 (Student’s t test between hUHRF1(PI5P) and hUHRF1(PI3P) experiments). Dashed line reflects full deuteration level of the corresponding peptide. Error bars represent SD from three independent measurements. **f**, Combined model of the crystal structure of the TTD-PHD module (4GY5) and the SRA domain (3CLZ) of hUHRF1, highlighting regions of interest and colored by domain. The numbered regions highlighted in blue (I, II and III) are indicating regions of slow HDX. Region IV highlighted in red shows fast HDX. Region (V) is highlighted in blue based on slow HDX as shown in Extended Data Fig. 6. The dotted lines point out the arbitrary continuation of the structure. **g**, Titration series of di-C 16:0 PI5P with fluorescently labeled protein domains (TTD and PHD) and Linker 2 of hUHRF1 analyzed by microscale thermophoresis. Data are plotted as average of three independent experiments; error bars correspond to SD. K_D_ values are listed in Extended Data Table 1. **h**, Titration series of wild type and mutated Linker 2 with fluorescently labeled wild type and mutated Linker 4 were analyzed in absence and presence of di-C 16:0 PI5P or PI3P by microscale thermophoresis. Data are plotted as average of three independent experiments; error bars correspond to SD. K_D_ values are listed in Extended Data Table 1.

Region I maps to the TTD surface groove (aka acidic patch or R pocket)^3,26^ that shares binding interface with either Linker 4/ PBR^3,27^ or with Linker 2^41^. Like Linker 4, Linker 2 is enriched in positively charged amino acids (RRK motifs, see Fig. 1). Among these residues R296 is crucial for interaction with the TTD surface groove^41^. For hUHRF1(PI5P) the peptides in the TTD surface groove showed 100 times delay to achieve the same level of deuteration as observed for hUHRF1(PI3P) (Fig. 4b).

The HDX profile of Linker 2 (region II) indicated that in the state of hUHRF1(PI3P) two thirds of the region remained unfolded while maintaining a stably folded structure in the rest of the peptide (compare the measured deuterium uptake with the calculated full deuteration level, see ref. 42). In contrast, in the state of hUHRF1(PI5P), deuterium uptake occurred in three states: only a few amino acid residues showed rapid HDX by 10^1^ s, an intermediate level of exchange occurred for several more residues between 10^3^ s and 10^4^ s followed by a third set of residues with added stability that didn’t undergo HDX past 10^5^ s (Fig. 4c). This pattern implied the induction of more stable secondary structures in Linker 2 in the presence of PI5P compared to PI3P.

Region III identified in the analysis maps to the binding site of the N-terminus of the H3-tail^18,19.^ The corresponding peptides showed very weak deuteration compared to the control set throughout the HDX time course (Fig. 4d), indicating that this region is stably locked in a state facing the TTD-Linker 2.

Peptides that map to the SRA domain (476-512 aa; region IV) showed higher deuteration levels when comparing hUHRF1(PI5P) with hUHRF1(PI3P) suggesting that this region gets exposed to the surroundings due to conformational rearrangement caused by PI5P binding (Fig. 4a,e,f and Extended Data Fig. 7a). The HDX patterns were consistent with half of this region forming loop structures and the other half acquiring stable secondary structures in the presence of PI5P. In the state of hUHRF1(PI3P) the number of loop residues is decreased and the region assumes a relatively stable secondary structure (Fig. 4e).

To evaluate whether slow/ less deuteration in the identified regions of hUHRF1 is caused by direct interaction with PI5P, we analyzed binding of the phospholipid to isolated Linker 2, TTD and PHD. In MST experiments, none of these regions showed any interaction with PI5P indicating that other parts of the protein such as Linker 4 are required for eliciting the observed effects (Fig. 4g). Next, we tested interaction of these regions with PI5P in presence of Linker 4. PI3P served as control in these experiments. In line with our earlier work^3^, we found that the TTD interacted with the apo-Linker 4 but that this interaction was lost in presence of PI5P (Extended Data Fig. 8a). In contrast, no interaction of the PHD domain with Linker 4 was detected in the presence or absence of PI5P (Extended Data Fig. 8b). The results suggested that the protection of this region observed in HDXMS is most likely due to conformational rearrangements of hUHRF1 induced by PI5P-binding.

Using isolated peptides, we found that Linker 2 interacted with Linker 4 in presence of PI5P but not in its absence. (Fig. 4h and Extended Data Fig. 8c). This effect was highly specific, since PI3P that can bind Linker 4 (Fig. 4h and Extended Data Fig. 5b) did not have a similar effect. Mutagenesis of R649 in Linker 4 that is implied in PI5P-binding abolished the interaction with Linker 2. Similarly, mutagenesis of R296 within Linker 2 caused loss of PI5P mediated binding of Linker 4. Collectively, our results implied that Linker 4 is essential for establishing a composite mode of hUHRF1 PI5P interaction. Upon binding of PI5P Linker 4 is released from the surface of the TTD. PI5P then mediates interaction of Linker 4 with Linker 2. The resulting conformational change is accompanied by additional changes in the PHD (more buried) and SRA (less buried) regions.

### PI5P directs synergism of hUHRF1 TTD-PHD in H3K9me binding

To further understand the allosteric regulation of hUHRF1 by PI5P we analyzed phosphoinositide binding of a series of different mutants of the full-length protein and monitored the TTD-PHD dependent synergism by measuring their H3-tail binding behavior in apo-vs. PI5P-bound state. In the apo-state the PBR* mutant of multiple charged residues showed only 4-fold enhancement for H3K9me3-binding compared to the wild type protein (Fig. 5a). This finding was consistent with this mutant adopting a “TTD open” state by releasing Linker 4 from the TTD surface groove. No further enhancement of the interaction with H3K9me3 was seen in the presence of PI5P, which is in agreement with this mutant not interacting with PI5P (Fig. 3, 5e). The results further confirmed that TTD unblocking alone is not sufficient for establishing a multivalent, synergistic H3K9me3-binding mode of hUHRF1.

**Fig. 5:**
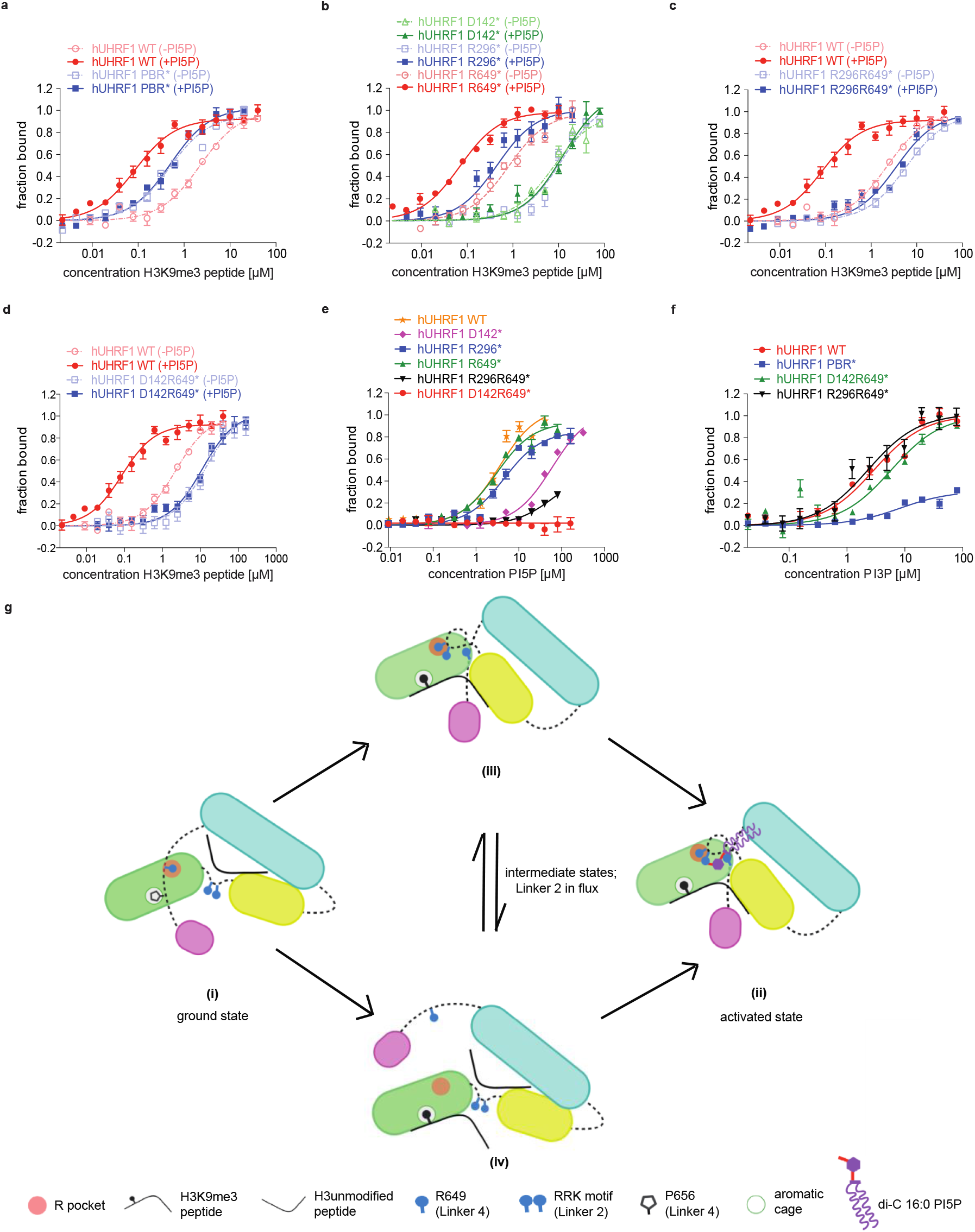
Mechanism of hUHRF1 allosteric activation by PI5P. **a** to **d**, Titration series of H3K9me3 peptide with fluorescently labeled, mutant, full-length hUHRF1 in absence and presence of di-C 16:0 PI5P were analyzed by microscale thermophoresis. PBR*: K644A, K646A, K648A, R649A, K650A, S651A; R296*: R296A; R649*: R649A; D142*: D142A; R296R649*: R296A, R649A; D142R649*: D142A, R649A. Data are plotted as average of three independent experiments; error bars correspond to SD. K_D_ values of all measurements are listed in Extended Data Table 1. **e**, Titration series of di-C 16:0 PI5P with fluorescently labeled hUHRF1 carrying the indicated mutations were analyzed by microscale thermophoresis. Data are plotted as average of three independent experiments; error bars correspond to SD. K_D_ values are listed in Extended Data Table 2. **f**, Titration series of di-C 16:0 PI3P with fluorescently labeled, WT and mutant, full-length hUHRF1 were analyzed by microscale thermophoresis. Data are plotted as average of three independent experiments; error bars correspond to SD. K_D_ values are listed in Extended Data Table 2. **g**, Putative conformational states of wild type and mutant hUHRF1 proteins. (i) In the apo/ ground state of wild type hUHRF1, the TTD domain is blocked by PBR/ Linker 4. Only the PHD domain is accessible for H3 N-terminal tail binding. (ii) PI5P recognizes and stabilizes a particular conformational state of hUHRF1. In this activated state, Linker 4 is displaced from the TTD surface; Linker 2 is positioned on the TTD surface groove and bridged to Linker 4 by PI5P. This conformational transition locks the TTD and PHD in a particular three-dimensional state that promotes multivalent, synergistic recognition of the H3K9me3 tail. In hUHRF1 PBR*, Linker 4 cannot block the TTD surface. Linker 2 remains dynamic and flexible allowing both compact (iii) and extended (iv) conformations. Despite free accessibility, it is not permanently bound to the TTD surface groove. In hUHRF1 R649* the TTD is unblocked. Like the wild type protein, it can interact with PI5P and form the activated conformational state for enhanced H3K9me3 recognition (ii). hUHRF1 R296* binds to PI5P only via the PBR/ Linker 4. While this releases the PBR-mediated TTD blocking, the mutant Linker 2 cannot be stably positioned on the TTD surface groove. In consequence hUHRF1 R296* is unable to adopt the activated conformation required for multivalent, synergistic H3K9me3 tail binding (iv). The conformational states of hUHRF1 R296R649* and hUHRF1 D142R649* are similar to that of hUHRF1 PBR* with the difference that PI5P is bound to the PBR. Linker 4 is displaced from the surface of the TTD but due to the mutations in the TTD surface groove or Linker 2 these cannot interact.

Although the R296* and R649* mutations abolished interaction with PI5P in isolated Linkers 2 and 4, these mutations had no effect on PI5P-binding in the context of the full-length protein (K_D_(R296*) = 5 μM, K_D_(R649*) = 3 μM) probably due to a mutation buffering effect (Fig. 5e and Extended Data Table 2)^43^. While the R649* mutation similar to the PBR* mutations unblocked the TTD inhibition, the unaffected PI5P-binding activated multivalent, synergistic H3K9me3 interaction of hUHRF1 with a binding strength similar to the wild type protein (K_D_ = 67 nM, Fig. 5b and Extended Data Table 1). In contrast, the R296* mutation showed no signs of TTD unblocking with its binding to H3K9m3 similar to wild type apo-hUHRF1 (K_D_ = 8.9 μM, Fig. 5b). While PI5P increased the interaction of this mutant with H3K9me3, the effect was significantly lower compared to the wild type protein (K_D_ = 0.4 μM, Fig. 5b). As the enhancement of binding was comparable to that of the R649* mutation in absence of PI5P, we deduced that the observed effect is due to the PI5P mediated release of Linker 4 from the TTD. But as Linker 2 R296* cannot access the TTD surface groove, no synergism between TTD and PHD is established.

To further test the hypothesis that PI5P induces TTD-PHD synergism via positioning Linker 2 in the surface groove of the TTD, we analyzed the D142* mutation that disrupts both Linker 2 and Linker 4 interactions with the TTD (i.e. the TTD is neither blocked by PBR/ Linker 4 nor activated by Linker 2)^3,27,41^. hUHRF1 D142* showed almost 20-fold weaker interaction with PI5P compared to the wild type protein (K_D_ = 60 μM, Fig. 5e). The results clearly implied that hUHRF1 must undergo three-dimensional rearrangements to acquire the PI5P-binding compatible active state. As expected, hUHRF1 D142* did not show any enhancement for H3K9me3 binding upon PI5P addition (Fig. 5b). To recapitulate a similar inactive state as D142*, we analyzed the R296R649* Linker 2 and Linker 4 double mutant and the TTD surface groove and Linker 4 double mutant D142R649*. Like D142*, these double mutants disrupt Linker 2 and Linker 4 interaction with the TTD surface groove. Importantly, both mutants did not bind to PI5P (Fig. 5e) and addition of the phospholipid did not enhance hUHRF1 H3K9me3-binding further, verifying that conformational rearrangements of hUHRF1 are necessary for both, PI5P interaction and induction of a multivalent, synergistic H3K9me3-binding mode (Fig. 5c,d).

Finally, we assessed the specificity of the allosteric interaction and conformational activation of hUHRF1 by measuring PI3P binding to the mutant proteins. PI3P binding to hUHRF1 was only disrupted by the PBR* but not the R296R649* and D142R649* mutations, indicating that PI5P but not PI3P is bound by an interface generated between TTD, Linker 2 and Linker 4 (Fig. 5f). Altogether, our findings are consistent with a model where PI5P binding via Linker 4 and Linker 2 releases Linker 4 from the TTD and simultaneously places Linker 2 into the TTD surface groove thereby establishing a conformational link between TTD and PHD that is highly favorable (i.e. synergistic) for H3K9me3-tail binding (Fig. 5g (ii)).

## Discussion

Here, we are reporting the first molecular details of allosteric regulation of a chromatin reader protein by a phosphoinositide. Our findings contrast and expand earlier work investigating isolated modules of hUHRF1 that while describing independent interaction modes did not reflect the regulation and multivalent, synergistic H3K9me3-binding functionality of the whole protein^28,44.^ We think the PI5P-induced transitions of hUHRF1 occur in two-steps: from a TTD blocked ground state (Fig. 5g (i)) through an TTD-open intermediate state (Fig. 5g (iii), (iv)) to a TTD-PHD synergistic activated state (Fig. 5g (ii)).

In the ground state the PBR/ Linker 4 is bound to the peptide-binding groove of the TTD (Fig. 5g (i)). R649 of Linker 4 is positioned into the R-pocket and especially interacts with residues D142 and E153 of the TTD surface^3,45^. Other PBR residues (K648, S651) also connect with the TTD surface strengthening the interaction. Finally, P656 of Linker 4 gets stably positioned into the aromatic cage of the TTD (Y191, F152 and Y188) that is required for K9me3 binding^45^. As consequence, the TTD surface is blocked for both, interaction with Linker 2 and also the H3 N-terminal tail.

The intermediate state of hUHRF1 is characterized by removal of the PBR/ Linker 4 from the TTD surface. Experimentally, it is obtained by PBR mutagenesis (PBR*) or binding to PI3P. In this state Linker 2 can access the TTD surface groove (Fig. 5g (iii)). But there is no stable association and Linker 2 positions fluctuate (Fig. 5g (iv)). Structural studies of isolated TTD-PHD have indeed shown that this dual module can adopt either a compact or extended confirmation depending on the position of Linker 2^28,44,46^. H3K9me3 binding either independently by the TTD (Linker 2 not bound by the TTD surface groove) or synergistically by the TTD and PHD (Linker 2 positioned in the surface groove) is possible in this state. In consequence, the intermediate state shows a slight preference for H3K9me3 over the unmodified H3-tail.

The activated state is characterized by strong synergism between the TTD and PHD domains (Fig. 5g (ii)). di-C 16:0 PI5P but not PI3P binding drives large conformational rearrangements that not only free the TTD from PBR/ Linker 4 inhibition, but also stabilizes the orientation of the PHD domain by firmly positioning Linker 2 on the TTD surface groove. This is accomplished by a composite interaction of Linker 4 and Linker 2 with PI5P that is not possible with the related PI3P. Indeed, mutagenesis of the TTD surface groove indicates that Linker 2 has to be on the TTD surface to be able to bind PI5P and to form a complex with PI5P-bound Linker 4 (Fig. 4h). Due to a favorable and stable orientation and distance of the TTD and PHD, multivalent, synergistic binding of H3K9me3 with the N-terminus of H3 bound by the PHD and the K9me3 moiety inserted into the aromatic cage of the TTD is enabled. Interestingly, this coupling of binding events is only mildly observed for methylated DNA ligase 1 (LIG1), another binding partner of UHRF1 targeting the TTD domain (Extended Data Fig. 8d)^47,48.^

Besides changes in TTD, Linker 2 and Linker 4, PI5P binding also affects the PHD and a region of the SRA domain (476-512 aa) that specifically recognizes methyldeoxycytosine bases in double stranded DNA (Fig. 4)^20,21^. Previous studies on isolated domains have shown that the SRA can bind to the PHD and partially inhibit its interaction with the unmodified N-terminus of the H3 tail^27^. We think that as a result of the PI5P-driven overall conformational rearrangements, the SRA-PHD interaction opens up which aids slight enhancement in unmodified H3-tail recognition (Fig. 1c and 2k). However, PI5P-binding did not affect the interaction of hUHRF1 with hemimethylated DNA (Extended Data Fig. 8e). As the SRA is crucial for the activation of the ubiquitin E3 ligase activity of hUHRF1^33^, it is tempting to speculate that the PI5P and hemimethylated DNA dependent conformations of the protein are exclusive to each other, thereby determining distinct functionalities of the protein.

Phospholipids are amphiphilic molecules that tend to form bicelles in aqueous solvent depending on the properties of the acyl chains. Binding analysis using PI5P bicelles (PI5P/DMPC/DHPC or PI5P/DHPC bicelles) vs. free PI5P showed that hUHRF1 prefers the non-bicelle state of PI5P (Extended Data Fig. 8f,g). Further considering the specificity (for the head group) and selectivity (for the acyl chain) for di-C 16:0 PI5P (Fig. 1a) and the effects of charge and hydrophopic properties changing mutations on PI5P binding (Fig. 3d,e), it seems that hUHRF1 recognizes individual molecule(s) of PI5P by interacting simultaneously with the head group and the acyl chains of the phospholipid. Determining the exact binding mode of PI5P will likely require refining existing structural methods for working with protein/ phospholipid complexes.

Several metabolites of energy homeostasis such as Acetyl-CoA, SAM, ATP, α-Ketoglutarate and NAD^+^ have been well investigated in their role as cosubstrates and cofactors of various chromatin-modifying enzymes^49,50^. Besides such direct effects, various other small cellular molecules and metabolites that exert more indirect chromatin regulation are emerging. As described here for hUHRF1/PI5P these often allosterically direct the functional state of chromatin factors. For instance, the class I histone deacetylase, HDAC3:SMRT complex structurally and functionally depends on inositol phosphate [Ins(1,4,5,6)P4]^51,52^. Further, autoinhibition of the chromatin remodeler ALC1/CHD1 is controlled by poly-ADP-ribose (PAR)^53^. *O*-acetyl-ADP-ribose (AAR) promotes association of multiple copies of Sir3 with Sir2/Sir4 and induces profound structural rearrangement in the SIR complex^54,55^. Given the complexities of chromatin control and the involved protein machinery, it is likely that more allosteric regulatory systems of chromatin factors and complexes will be discovered. Defining the physiological role of these regulation events remains a significant challenge, as the experimental tools for targeted manipulation of the small cellular molecules are difficult to establish. Indeed, up to date there are no methods for interfering specifically with the nuclear PI5P pool. The enzyme systems and putative signaling events controlling nuclear PI5P are complex and difficult to control^56-58^. Chemical biology approaches for derivatizing PI5P^59^ and nuclear injection^60,61^ coupled to single cell analysis might be promising avenues in this regard to deduce the functional role of UHRF1/ PI5P regulation.

## Methods

### Plasmids

All cloning was done using human UHRF1 cDNA (NM_001048201) and following standard procedures. For bacterial expression of recombinant proteins, the following constructs were used: pETM13 hUHRF1 (aa 1-793)-6xHIS, pETM13 Linker 4 (aa 605-675)-6xHIS, pETM13 RING (aa 675-793)-6xHIS, pETM13 UBL-TTD-PHD-SRA (aa 1-619)-6xHIS, pET16b 10xHIS-TEV-TTD (aa 126-285), pETM13 PHD (aa 301-376)-6xHIS and pET16b 10xHIS-TEV-USP7 UBL1-2 (aa 560-792). Point mutagenesis was performed following the QuickChange Protocol (Stratagene). Details of constructs used in this study are available upon request.

### Lipids

Phosphatidylinositol 5-phosphate diC16 (di-C 16:0 PI5P; Echelon #P5016); Phosphatidylinositol 4-phosphate diC16 (di-C 16:0 PI4P; Echelon #P4016); Phosphatidylinositol 3-phosphate diC16 (di-C 16:0 PI3P; Echelon #P3016); Phosphatidylinositol 4,5-bisphosphate diC16 (di-C 16:0 PI(4,5)P2; Echelon #P4516); Phosphatidylinositol 3,5-bisphosphate diC16 (di-C 16:0 PI(3,5)P2; Echelon #P3516); Phosphatidylinositol 5-phosphate diC8 (di-C 8:0 PI5P; Echelon #P-5008); Inositol 1,5-bisphosphate (Ins(1,5)P2; Echelon #Q-0015); 1,2-dioleoyl-sn-glycero-3-phospho-(1’-myo-inositol-5’-phosphate) (di-C 18:1 PI5P; Avanti Polar Lipids #850152P); 1-heptadecanoyl-2-(5Z,8Z,11Z,14Z-eicosatetraenoyl)-sn-glycero-3-phospho-(1’-myo-inositol-5’-phosphate) (17:0,20:4 PI5P; Avanti Polar Lipids #LM1902); 1,2-dipalmitoyl-sn-glycero-3-phosphoethanolamine-N-(biotinyl) (sodium salt) (di-C 16:0 Biotinyl PE; Avanti Polar Lipids #870285); 1,2-dimyristoyl-sn-glycero-3-phosphoethanolamine (di-C 14:0 PE; Avanti Polar Lipids #850745); 1,2-dioleoyl-sn-glycero-3-phosphoethanolamine-N-(biotinyl) (sodium salt) (di-C 18:1 Biotinyl PE; Avanti Polar Lipids #870282); 1,2-distearoyl-sn-glycero-3-phosphoethanolamine-N-[methoxy(polyethylene glycol)-550] (ammonium salt) (di-C 18:0 PEG550PE; Avanti Polar Lipids #880520); Cholesterol-(polyethylene glycol-600) (Avanti Polar Lipids #880001); 1,2-dipalmitoyl-sn-glycero-3-phosphate (di-C 16:0 PA; Avanti Polar Lipids #830855); 1,2-Dimyristoyl-sn-Glycero-3-Phosphocholine (DMPC; Anatrace #D514); 1,2-Diheptanoyl-sn-Glycero-3-Phosphocholine (DHPC; Anatrace #D607).

### Peptides

All peptides were custom synthesized and purified to >90% by Synpeptide Co: H3 unmodified (1-20 aa) = ARTKQTARKSTGGKAPRKQL-K(biotin); H3K9me3 (1-20 aa) = ARTKQTARK(me3)STGGKAPRKQL-K(biotin); human LIG1 (118-128 aa) = IPKRRTARKQL-K(biotin); LIG1me3 (118-128 aa) = IPKRRTARK(me3)QL-K(biotin); hUHRF1 Linker 2 (286-306 aa) = GSPMVDNPMRRKSGPSCKHC-K(biotin); hUHRF1 Linker 2 (286-306 aa) R296A = GSPMVDNPMRAKSGPSCKHC-K(biotin).

### Protein expression and purification

Proteins were expressed in BL21-DE3 RIL *E*.*coli* cells. Bacterial cultures growing at 30°C with shaking were induced with 0.75 mM IPTG at OD_600_ 0.5-0.6. After continued growth at 25°C for 3 hours, cells were harvested and lysed. His-tagged proteins were purified on HisPur Ni-NTA resin (Thermo Fischer) according to the manufacturer’s protocols. Eluates were dialyzed against 50 mM Tris-HCl (pH 8.0), 150 mM NaCl, 10% v/v glycerol and 1 mM DTT. Proteins were stored at 4°C until further use.

### Microscale thermophoresis

C-terminally His6-tagged proteins were labeled using the Monolith His-tag labeling kit RED-tris-NTA (Nanotemper #MO-L008) following the manufacturer’s protocol. Briefly, 400 nM protein was incubated with 100 nM of His-tag labeling dye in MST buffer (20 mM HEPES-NaOH (pH 7.9), 150 mM NaCl, 0.05% v/v Tween-20) for 30 minutes at room temperature. Labeled proteins were cleared by centrifugation at 15000 x g for 10 minutes at 4°C.

Fluorophore-labeled protein was incubated with various ligands at room temperature for 15 minutes before measuring on Monolith NT.115 (NanoTemper, 80% LED power, 40% MST power). For competition assays, protein-DNA or protein-PIP concentrations were kept constant. Data points were fitted using the following equation:

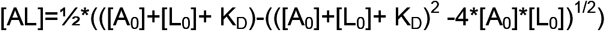

K_D,_ dissociation constant; [A_0_], concentration of fluorescent molecule; [L_0_], concentration of ligand/ binding partner; [AL], concentration of the complex of A and L. All binding measurements were performed in triplicate as biological and technical replicates with at least two independent protein preparations.

### Peptide pull-down

40 μl streptavidin paramagnetic beads (MagneSphere, Promega) were washed three times with PBS in low-binding tubes. 10 μg biotin-labeled hUHRF1 Linker 2 peptides or water was added for 1 hr at room temperature with rotation followed by three washes with PBS. 10 μg recombinant hUHRF1 Linker 4 peptide in 500 μL binding buffer (20 mM HEPES-NaOH (pH 7.9), 150 mM NaCl, 0.05% v/v Tween-20) were added and reactions were incubated for 1 hour at room temperature with rotation in absence and presence of 10-fold molar excess of di-C 16:0 PI5P or di-C 16:0 PI3P (final concentrations 2.15 μM Linker 4 and 21.5 μM PIP). Beads were washed three times with binding buffer and recovered material was eluted in 30 μL PDelute buffer (50 mM Tris-HCl (pH 8.0), 25% v/v glycerol, 0.25% w/v bromophenol blue, 1 mM EDTA, 2% w/v SDS, 1 mM TCEP) by boiling for 5 min.

### HDXMS measurements and analysis

hUHRF1 and H3 peptides or hUHRF1(PI5P/ PI3P) were mixed at final concentrations of 1 μM, 10 μM and 16.8 μM in buffer containing 50 mM Tris-HCl (pH 7.9), 150 mM NaCl, 1 mM DTT and incubated for 30 min at room temperature to allow complex formation. Deuterium on-exchange was carried out at 4**°**C by mixing 5 μL of each sample with 15 μL of deuterium on-exchange buffer (10 mM Tris-HCl (pD 8.3), 150 mM NaCl, in D_2_O) yielding a final D_2_O concentration of 75%. pD values for deuterium-based buffers were calculated as pD = pH + 0.4138. Upon mixing with on-exchange buffer, the amide protons are replaced over time with deuterons yielding an increased peptide mass. To quench the deuterium exchange reaction, samples (20 μL) were mixed with 30 μl of ice-cold quench buffer (500 mM guanidine hydrochloride, 10% v/v glycerol, and 0.8% *v/v* formic acid, for a final pH of 2.4). Samples were rapidly frozen in liquid nitrogen and stored at -80**°**C until further use.

For MS analysis, each sample (50 μL) was thawed on ice and loaded onto an in-house packed pepsin column for digestion. To this end, pepsin (Sigma) was coupled to POROS 20 AL support (Applied Biosystems), and the immobilized pepsin was packed into a column housing (2 mm × 2 cm, Upchurch). Pepsin-digested peptides were captured on a TARGA C8 5 μm Piccolo HPLC column (1.0 × 5.0 mm, Higgins Analytical) and eluted through an analytical C18 HPLC column (0.3 × 75 mm, Agilent) by a shaped 12–100% buffer B gradient at 6 μL/min (Buffer A: 0.1% formic acid; Buffer B: 0.1% formic acid, 99.9% acetonitrile). Eluting peptides were electrosprayed into the mass spectrometers (Exactive Plus EMR-Orbritrap or Q-exactive HF, both Thermo Fisher Scientific). Data were recorded using Xcalibur software (Thermo Fisher Scientific). MS only data were acquired for the time course analysis. MS/MS data were collected in data-dependent mode to sequence hUHRF1 peptides resulting from pepsin digestion. Peptides were identified by database searching using SEQUEST (Bioworks v3.3.1 and Proteome Discoverer V 2.4, Thermo Fisher Scientific) with a peptide tolerance of 8 ppm and a fragment tolerance of 0.1 AMU.

A MATLAB based program, ExMS2 was used to prepare the pool of peptides based on SEQUEST output files. EXMS2 (for hUHRF1-H3 peptides +/-PI5P measurements) and HDExaminer software (for hUHRF1-PI5P/ PI3P measurements) were used to process and analyze the HDXMS data. The software identifies the peptide envelope centroid values for non-deuterated as well as deuterated peptides and uses the information to calculate the level of peptide deuteration for each peptide at each timepoint. Each individual deuterated peptide was corrected for loss of deuterium label during HDXMS data collection by normalizing to the maximal deuteration level of that peptide (measured in a “fully deuterated” (FD) reference sample). The FD sample was prepared in 75% deuterium to mimic the exchange experiment but under acidic denaturing conditions (0.5% formic acid), and incubated over 48 hours to allow each amide proton position along the entire polypeptide to undergo full exchange. The software automatically performs the correction when provided with the FD file. For each peptide, we compared the extent of deuteration as measured in the FD sample to the theoretical maximal deuteration (i.e., if no back-exchange occurs). The median extent of back-exchange in our datasets was 15-20%.

### NMR spectroscopy

All NMR spectra were acquired on Bruker Avance NEO NMR spectrometers operating at 700 and 950 MHz equipped with sensitive triple resonance TCI cryoprobes. NMR samples for the backbone assignment of 200 µM uniformly ^15^N,^13^C-double labeled Linker 4 (hUHRF1 aa 605-675)-6xHIS) were prepared in 50 mM Tris-HCl (pH 6.5), 150 mM NaCl, 90%/10% *v/v* H_2_O/D_2_O, and 1 mM DTT. Complete sequence specific backbone resonance assignment of Linker 4 was obtained at 15°C from 3D triple-resonance HSQC-based HNCACB, CBCA(CO)NH, HNCA, HN(CO)CA, HN(CA)CO and HNCO spectra (https://pubs.acs.org/doi/abs/10.1021/bi00471a022). Spectra were processed in Topspin 4.0.7 and analyzed in CARA (http://www.cara.nmr-software.org/). The side-chain chemical shifts were assigned based on the following experiments: heteronuclear 2D ^1^H-^13^C HSQC and 3D C(CO)NH, 3D (H)CCH-TOCSY and H(C)CH-TOCSY (mixing times 16.2 ms) and 3D ^13^ C-edited HSQC-NOESY (mixing time 100 ms) covering the aliphatic region. The titrations with phospholipids were monitored with 2D ^1^H-^13^C HSQC spectra at 25°C recorded on 20 µM uniformly ^15^N,^13^C-labeled Linker 4 in 50 mM Tris-HCl (pH 7.9), 150 mM NaCl 1 mM DTT and PI5P/PI4P in 1:10 ratio.

### Bicelle formation and analysis

di-C 16:0 PI5P/DMPC/DHPC and di-C 16:0 PI5P/DHPC bicelles at 0.4 molar ratio of lipids were produced as described^62^. DMPC, DHPC and di-C 16:0 PI5P were dissolved in water. The slurries were vortexed and sonicated until the material was fully dissolved. di-C 16:0 PI5P (250 µM), DMPC (250 µM) and DHPC (1 mM) were mixed to obtain a sample with a total lipid ([PC] ≡ [PI5P+DMPC] + [DHPC]) concentration of 1.5 mM. di-C 16:0 PI5P/DHPC bicelles were prepared by mixing 500 µM of PI5P with 1 mM DHPC. This mixture was subjected to several cycles of gentle heating to 42 °C combined with vortexing until a clear non-viscous solution was obtained. Bicelles were incubated on ice for 15 minutes in 20 mM HEPES-NaOH (pH 7.9), 150 mM NaCl at a concentration of 150 µM. 4 µL of sample were added to a continuous carbon grid after glow discharge (Solarus, Gatan). After 1 minute incubation, samples were blotted with filter paper and stained three times with 2% *w/v* uranyl formate. The stain was removed by blotting with filter paper and the grids were dried at room temperature before imaging on a Thermo Fisher Scientific Tecnai Twin microscope (Gatan UltraScan 4000, 120 keV with a 4k x 4k CCD camera).

## Acknowledgement

We thank Abrar Aljahani for help with MST assays, Dr. Sarah Kreuz for valuable scientific input in the course of this study and members of the Fischle laboratory for stimulating discussions. This work was supported by the King Abdullah University of Science and Technology (intramural funds and award OSR-2015-CRG-2616 of the KAUST Office of Sponsored Research to W.F.) and NIH grants R35GM130302 (to B.E.B.) and F32GM128265 (to L.Z.).

## Author Contributions

P.M. and W.F. conceived and designed the project. P.M. performed all sample preparation and did the majority of the quantitative measurements. Z.Y. performed some binding assays and provided negative staining of lipid bicelles. V.K. measured and analyzed NMR spectra. L.J. directed the NMR part of the work. B.E.B. designed, contributed to and supervised HDXMS experiments. P.M., L.Z. and D.B. performed HDXMS measurements. P.M. analyzed the HDXMS data. W.F. supervised the course of the study. P.M. and W.F. wrote the manuscript. All authors discussed the results and commented on the manuscript.

