## Supplementary for "Molecular basis of UHRF1 allosteric activation for synergistic histone modification binding by PI5P"

[1]Biological and Environmental Science and Engineering Division, King Abdullah University of Science and Technology, Thuwal 23955, Saudi Arabia. [2]Department of Biochemistry and Biophysics, Penn Center for Genome Integrity, Epigenetics Institute, Perelman School of Medicine, University of Pennsylvania, Philadelphia, PA 19104, USA. [3]Core Laboratories, King Abdullah University of Science and Technology, Thuwal 23955, Saudi Arabia.

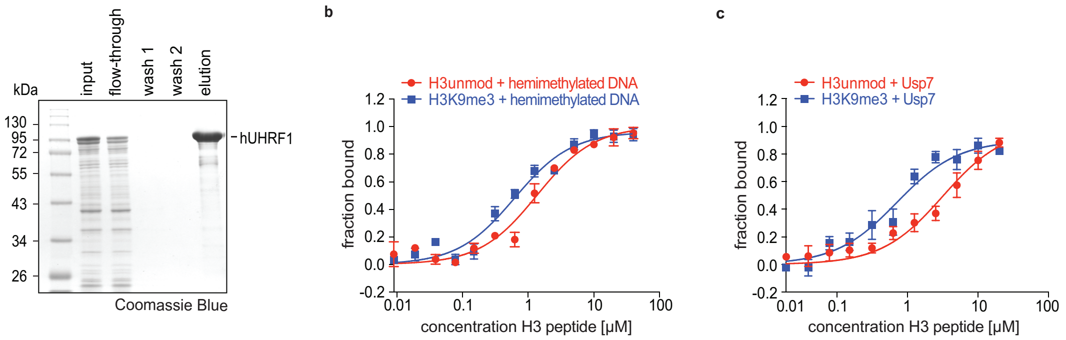


**Extended Data Fig. 1: Purification of full-length hUHRF1 and interaction with different ligands.** (**a**) His-tag affinity purification of full-length recombinant hUHRF1. Samples were run on SDS-PAGE and stained with Coomassie Blue. (**b** and **c**) Titration series of the indicated H3 peptides with fluorescently labeled recombinant hUHRF1 and in presence of hemimethylated DNA (b) or the UBL1/2 domain of hUSP7 (c) both at 60% saturation level were analyzed by microscale thermophoresis. Data are plotted as average of three independent experiments; error bars correspond to SD. $K_D$ values of all measurements are listed in Extended Data Table 1.

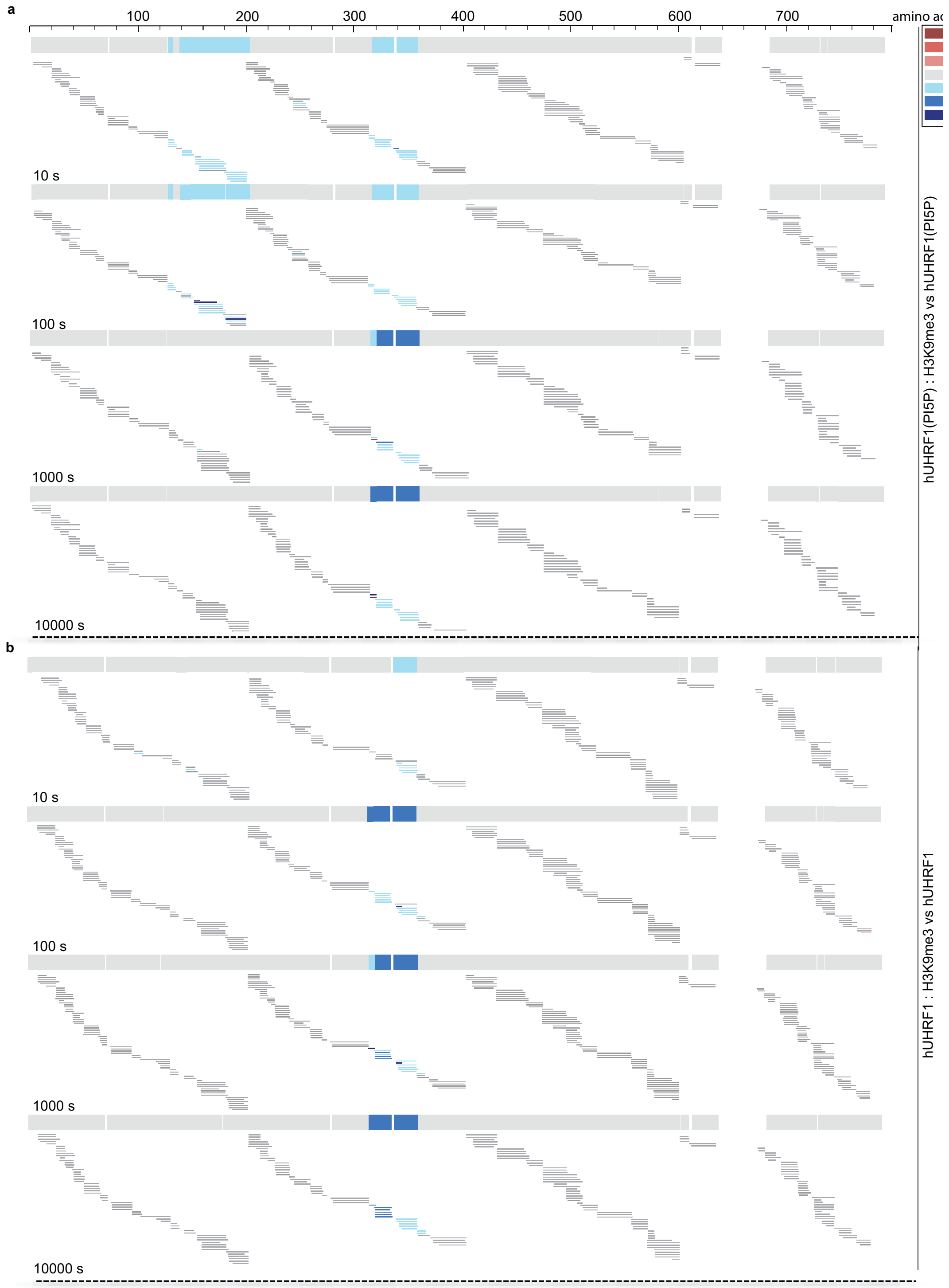


**Extended Data Fig. 2: Peptide data corresponding to the HDX consensus plots of Fig. 2a for hUHRF1 interacting with H3K9me3.**
(**a** and **b**) Percent difference in deuteration for each peptide (i.e. HDX), represented by horizontal bars was calculated between hUHRF1(PI5P) complex (a) and apo-hUHRF1 (b) in presence or absence of an H3K9me3 peptide at four time points (10 s – 10000 s). Peptides exhibiting increased protection from deuteration upon binding to the H3K9me3 peptide are shown in blue (i.e. slower HDX), whereas those with no change in deuteration uptake are shown in grey. For each time point the consensus pattern at each hUHRF1 residue is displayed on top of the peptide plots.

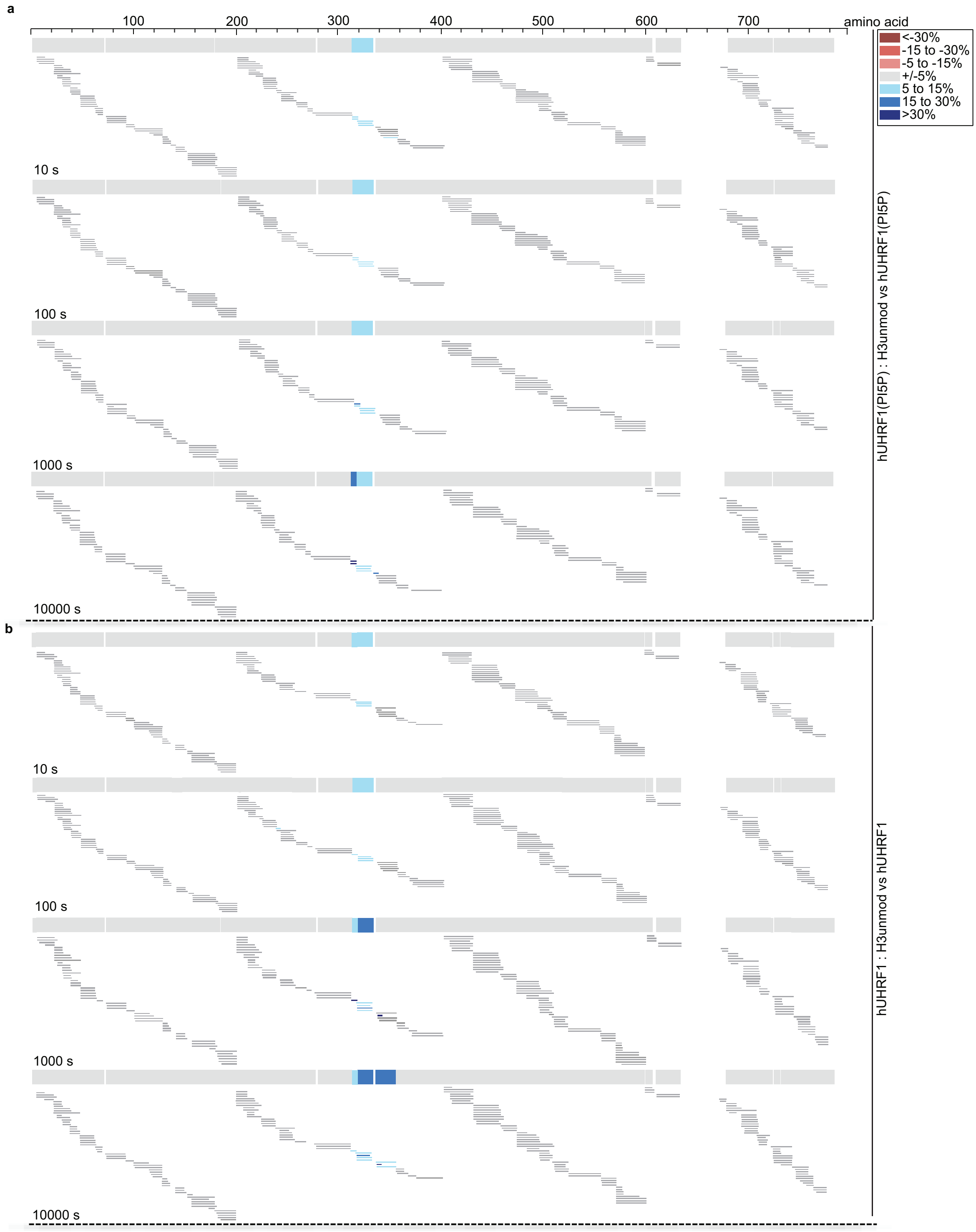


**Extended Data Fig. 3: Peptide data corresponding to the HDX consensus plots of Fig. 2a for hUHRF1 interacting with H3unmodified.**

(**a** and **b**) Percent difference in deuteration for each peptide (i.e. HDX), represented by horizontal bars was calculated between hUHRF1(PI5P) complex (a) and apo-hUHRF1 (b) in presence or absence of an H3unmodified peptide at four time points (10 s – 10000 s). Peptides exhibiting increased protection from deuteration upon binding to the H3unmodified peptide are shown in blue (i.e. slower HDX), whereas those with no change in deuteration uptake are shown in grey. For each time point the consensus pattern at each hUHRF1 residue is displayed on top of the peptide plots.

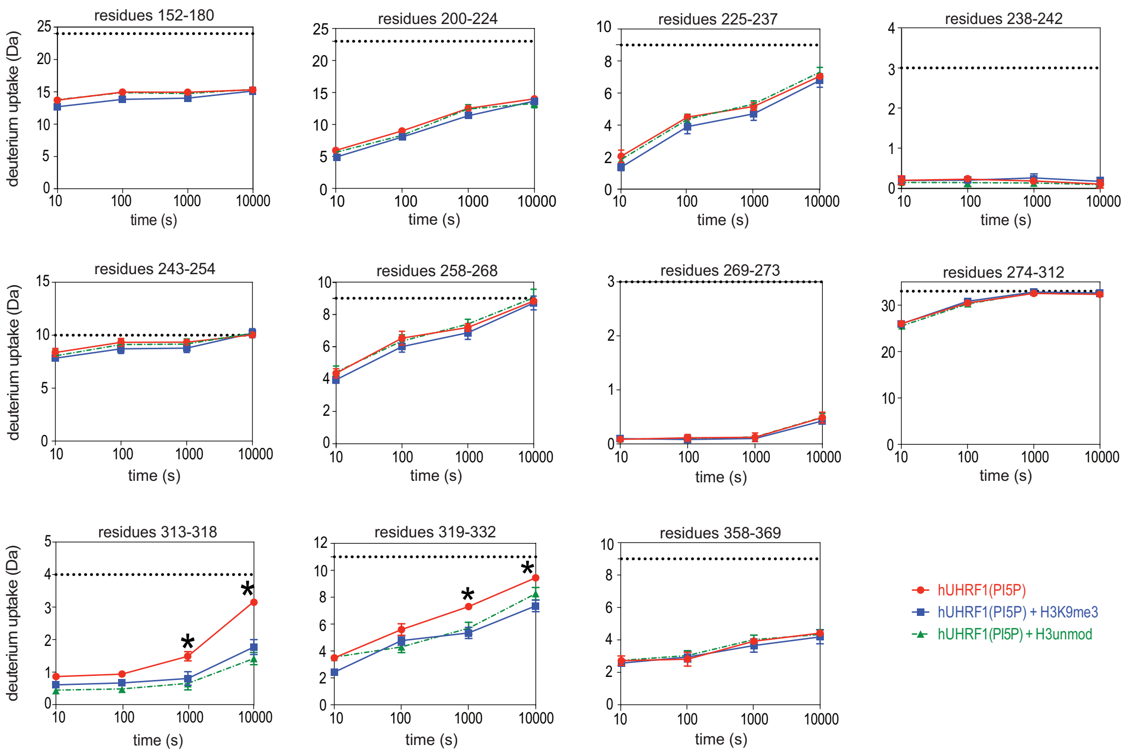


**Extended Data Fig. 4: Peptide deuteration curves for representative peptides covering the region from TTD to PHD of hUHRF1.**
Time curves are shown for the hUHRF1(PI5P) complex alone, or in presence of the indicated H3 peptides. The regions of hUHRF1 covered by the different identified peptides are listed on top of the graphs. Each data point reflects the average of three measurements. The level of deuteration was adjusted relative to the fully deuterated (FD) sample (see Materials and Methods section for details). Error bars correspond to SD from three independent measurements. The asterisks mark time points with $P < 0.01$ (Student's t test between hUHRF1(PI5P) and hUHRF1(PI5P)/H3K9me3 experiments).

a

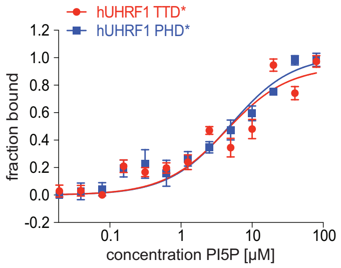


b

| lipid ligands | Linker 4 [µM] |
|---|---|
| 16:0 PI5P | 2.8 ± 0.4 |
| 16:0 PI4P | NB |
| 16:0 PI3P | 5.8 ± 0.2 |
| 16:0 PI3,5P2 | NB |
| 16:0 PI4,5P2 | NB |
| 18:1 PI5P | NB |
| 17:0 20:4 PI5P | NB |
| 8:0 PI5P | NB |
| 16:0 PE | NB |
| 18:1 PE | NB |
| cholesterol | NB |
| 1,5 IP | NB |
| 16:0 PA | NB |

**Extended Data Fig. 5: Interaction of different regions of hUHRF1 with phospholipids.**
(**a**) Titration series of di-C 16:0 PI5P with fluorescently labeled recombinant, mutant hUHRF1 were analyzed by microscale thermophoresis. TTD*: Y188A, Y191H; PHD*: D334, D337A. Data are plotted as average of three independent experiments; error bars correspond to SD.
(**b**) Binding of Linker 4 of hUHRF1 to phospholipids and other lipids of the indicated composition was analyzed by microscale thermophoresis. $K_D$ deduced as average from three independent titration measurements is listed. Error corresponds to SD. NB, not binding. $K_D$ values are listed in Extended Data Table 1.

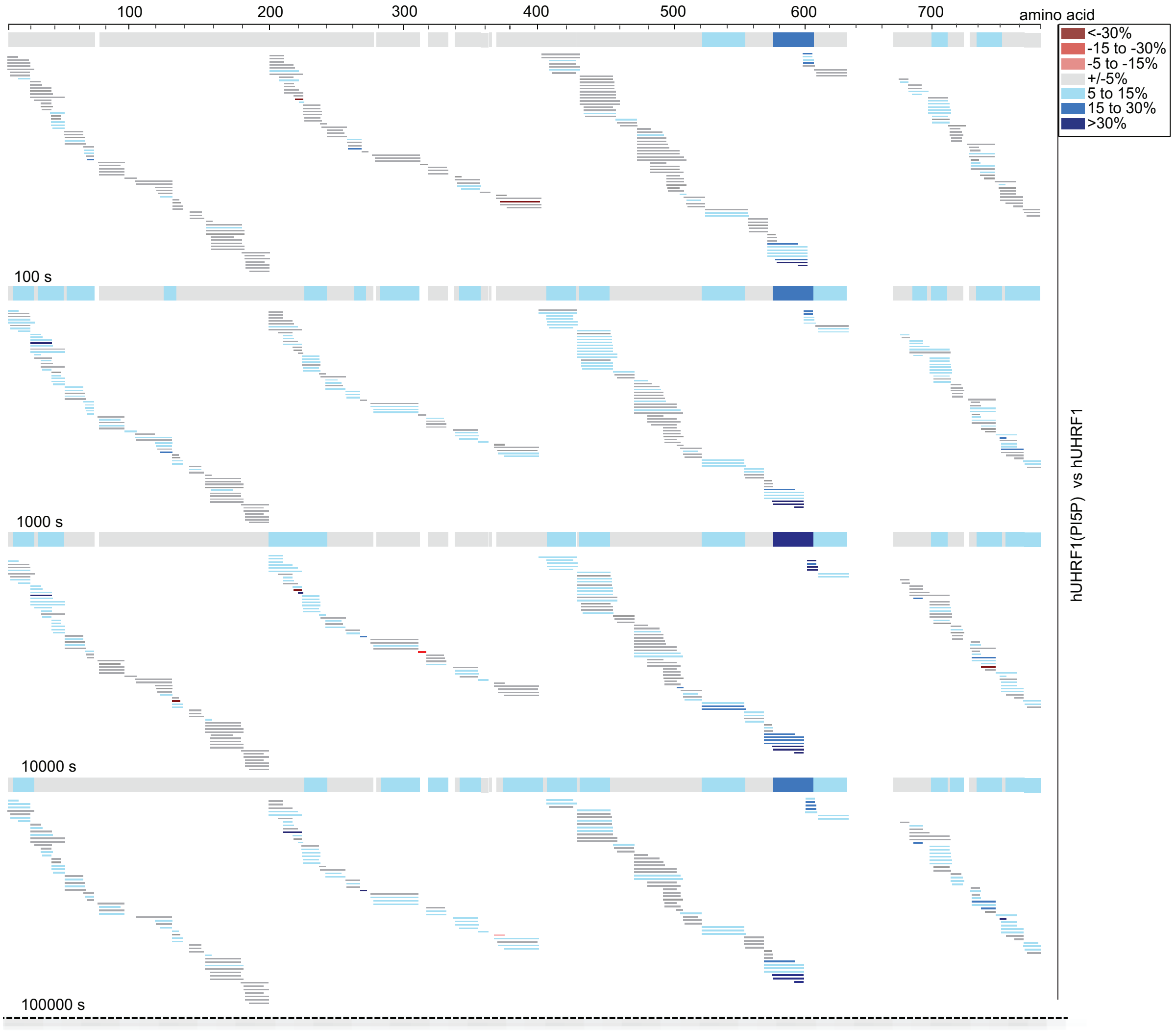


**Extended Data Fig. 6: Peptide data corresponding to the HDX consensus plots of Fig. 4a for hUHRF1 interacting with PI5P.**
Percent difference in deuteration for each peptide (i.e. HDX), represented by horizontal bars was calculated between hUHRF1(PI5P) complex and apo-hUHRF1 at four time points (10 s – 10000 s). Peptides exhibiting increased protection from deuteration upon binding to di-C 16:0 PI5P are shown in blue (i.e. slower HDX), whereas those with no change in deuteration uptake are shown in grey.

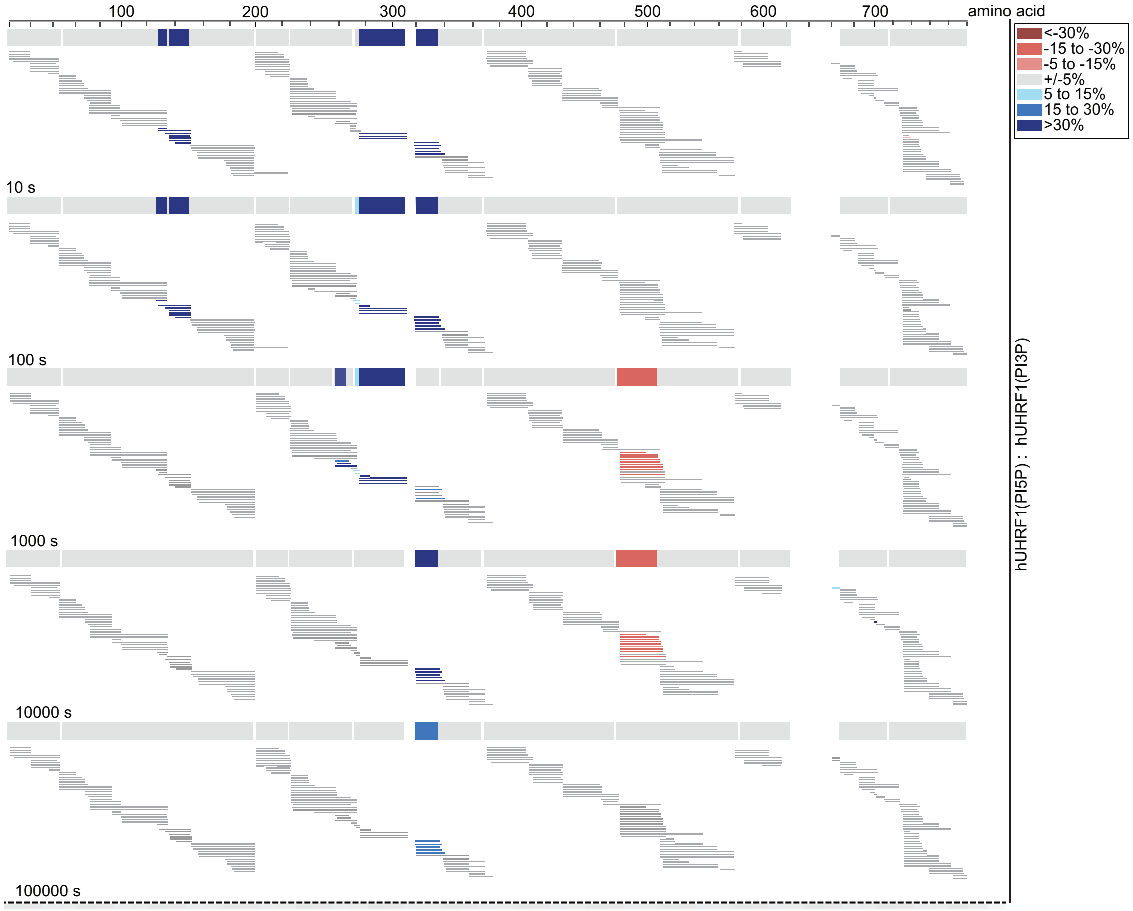


**Extended Data Fig. 7: Peptide data corresponding to the HDX consensus plots of Fig. 4a for hUHRF1 interacting with PI5P vs. PI3P.**

Percent difference in deuteration for each peptide (i.e. HDX), represented by horizontal bars was calculated between hUHRF1(PI5P) and hUHRF1(PI3P) complexes at four time points (10 s – 10000 s). Peptides exhibiting increased protection from deuteration upon binding to di-C 16:0 PI5P are shown in blue (i.e. slower HDX), whereas those with decreased protection from deuteration are shown in red (i.e. faster HDX). Peptides with no change in deuteration uptake between the two complexes are shown in grey.

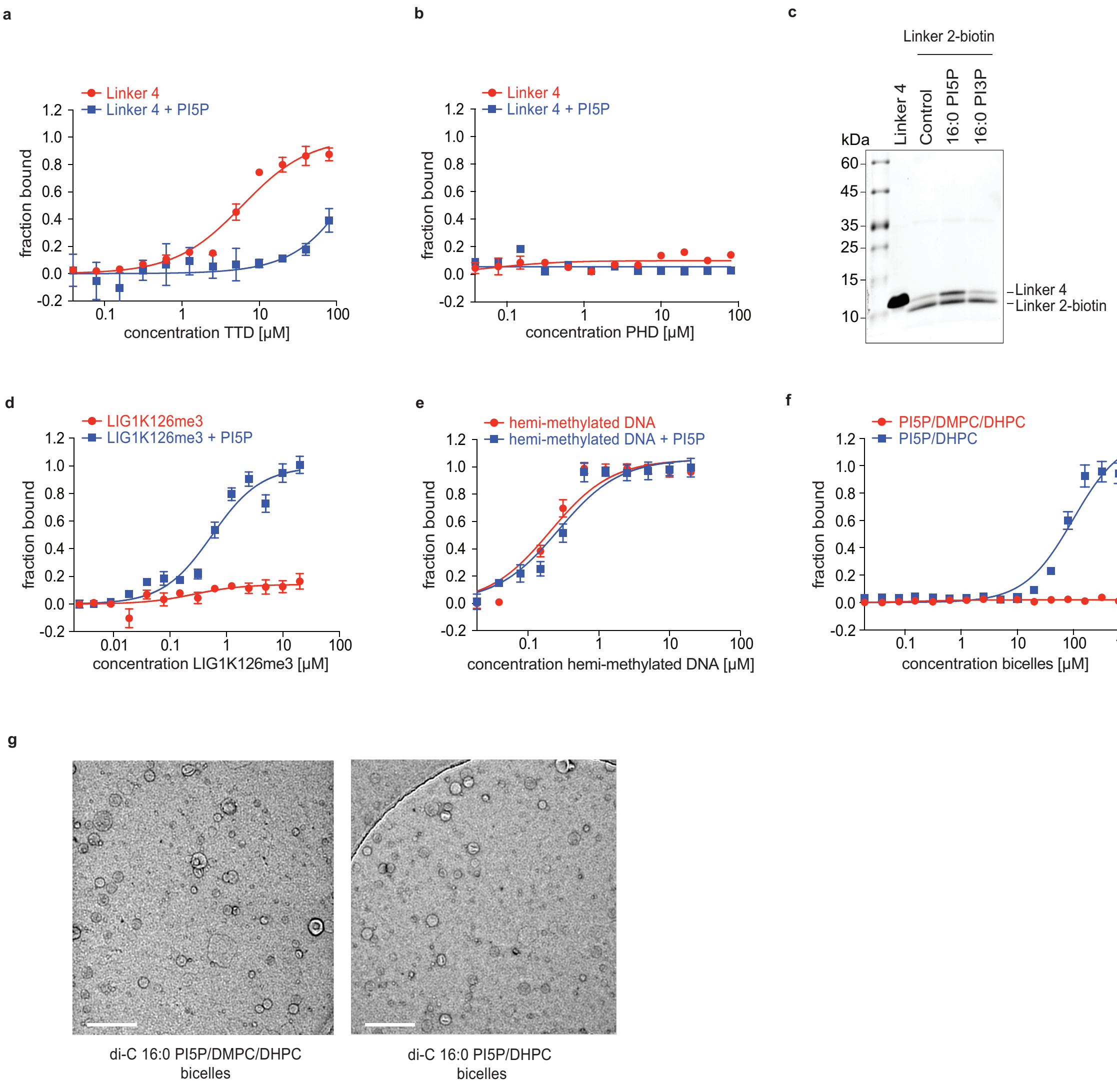


**Extended Data Fig. 8: Binding analysis of hUHRF1 and its domains with different ligands and in different physical state.**

(**a** and **b**). Titration series of the isolated, recombinant TTD (a) and PHD (b) domains of hUHRF1 with fluorescently labeled recombinant Linker 4 were analyzed in absence and presence of di-C 16:0 PI5P by microscale thermophoresis. Data are plotted as average of three independent experiments; error bars correspond to SD. $K_D$ values are listed in Extended Data Table 1.

(**c**) A biotinylated peptide corresponding to Linker 2 of hUHRF1 was used in pull-down experiments with recombinant Linker 4 of hUHRF1 in absence and presence of di-C 16:0 PI5P and di-C 16:0 PI3P. Recovered material was analyzed by SDS PAGE gel and stained with Coomassie. Running positions of molecular weight markers (MW), Linker 2 and Linker 4 are indicated.

(**d** and **e**). Titration series of tri-methylated LIG1 peptide (LIG1K126me3) (d) and hemi-methylated DNA (e) with fluorescently labeled recombinant hUHRF1 were analyzed in absence and presence of di-C 16:0 PI5P by microscale thermophoresis. Data are plotted as average of three independent experiments; error bars correspond to SD. $K_D$ values are listed in Extended Data Table 1.

(**f**) Titration series of lipid bicelles of the indicated composition with fluorescently labeled recombinant hUHRF1 were analyzed by microscale thermophoresis. Two different types of di-C 16:0 PI5P embedded bicelles were used. In the PI5P/DMPC/DHPC bicelles the concentration of PI5P was lower compared to the PI5P/DHPC bicelles, where the DMPC molecules were replaced by PI5P. Data are plotted as average of thre independent experiments; error bars correspond to SD. $K_D$ values are listed in Extended Data Table 2.

(**g**) Lipid bicelles formed by PI5P/DMPC/DHPC and PI5P/DHPC were analyzed by electron microscopy using negative staining. Scale bar, 200 nm.

**Extended Data Table 1: Dissociation constants ($K_D$ in µM) of interaction of different complexes of wild type or mutant hUHRF1 with different ligands as determined by microscale thermophoresis.**

hUHRF1 TTD*: Y188A, Y191H; hUHRF1 PHD*: D334, D337A; hUHRF1 PBR*: K644A, K646A, K648A, R649A, K650A, S651A; hUHRF1 R296R649*: R296A, R649A; hUHRF1 R649*: R649A; hUHRF1 R296*: R296A; hUHRF1 D142*: D142A; hUHRF1 D142R649*: D142A, R649A; Linker 4 R649*: hUHRF1 (aa 605-675) R649A; Linker 4 R296*: hUHRF1 (aa 605-675) R296A. NB, not binding; – not determined.

<table>
<tr><th rowspan="2"></th><th colspan="2">apo form</th><th colspan="2">ligand bound form</th></tr>
<tr><th>H3unmod</th><th>H3K9me3</th><th>H3unmod</th><th>H3K9me3</th></tr>
<tr><td>hUHRF1 (PI5P)</td><td rowspan="6">2.9 +/- 0.4</td><td rowspan="6">2.2 +/- 0.3</td><td>1.2 +/- 0.2</td><td>0.08 +/- 0.01</td></tr>
<tr><td>hUHRF1 (PI3P)</td><td>1.4 +/- 0.2</td><td>0.65 +/- 0.1</td></tr>
<tr><td>hUHRF1 (PE)</td><td>2.7 +/- 0.3</td><td>0.9 +/- 0.1</td></tr>
<tr><td>hUHRF1 (PI4P)</td><td>0.6 +/- 0.08</td><td>0.7 +/- 0.1</td></tr>
<tr><td>hUHRF1 (hemi-methylated DNA)</td><td>1.3 +/- 0.2</td><td>0.6 +/- 0.1</td></tr>
<tr><td>hUHRF1 (Usp7)</td><td>3.1 +/- 0.6</td><td>0.7 +/- 0.1</td></tr>
<tr><td>hUHRF1 TTD* (PI5P)</td><td>5 +/- 0.8</td><td>6.7 +/- 1</td><td>2.1 +/- 0.3</td><td>1.3 +/- 0.2</td></tr>
<tr><td>hUHRF1 PHD* (PI5P)</td><td>NB</td><td>NB</td><td>NB</td><td>6.4 +/- 2</td></tr>
<tr><td>hUHRF1 PBR* (PI5P)</td><td>2.6 +/- 0.4</td><td>0.5 +/- 0.08</td><td>2.7 +/- 0.6</td><td>0.5 +/- 0.06</td></tr>
<tr><td>hUHRF1 R296R649* (PI5P)</td><td>6.1 +/- 0.8</td><td>5.7 +/- 0.4</td><td>3.7 +/- 0.9</td><td>3.5 +/- 0.6</td></tr>
<tr><td>hUHRF1 R649* (PI5P)</td><td>5.8 +/- 2.2</td><td>0.7 +/- 0.1</td><td>4.9 +/- 1</td><td>0.067 +/- 0.01</td></tr>
<tr><td>hUHRF1 R296* (PI5P)</td><td>8.3 +/- 0.7</td><td>8.9 +/- 0.8</td><td>1.2 +/- 0.1</td><td>0.4 +/- 0.08</td></tr>
<tr><td>hUHRF1 D142* (PI5P)</td><td>–</td><td>8.4 +/- 2</td><td>–</td><td>9.2 +/- 2.1</td></tr>
<tr><td>hUHRF1 D142R649* (PI5P)</td><td>–</td><td>13.4 +/- 2.8</td><td>–</td><td>11.4 +/- 2.3</td></tr>
</table>

| | LIG1K126me3 | LIG1K126me3 + PI5P | hemimethylated DNA | hemimethylated DNA + PI5P |
|---|---|---|---|---|
| hUHRF1 | NB | 0.6 +/- 0.1 | 0.2 +/- 0.05 | 0.3 +/- 0.06 |

<table>
<tr><th rowspan="2"></th><th colspan="4">apo form</th><th colspan="4">ligand bound form</th></tr>
<tr><th>TTD</th><th>PHD</th><th>Linker 2</th><th>Linker 4</th><th>TTD</th><th>PHD</th><th>Linker 2</th><th>Linker 4</th></tr>
<tr><td>Linker 4 (PI5P)</td><td rowspan="2">6 +/- 1.2</td><td rowspan="2">NB</td><td rowspan="2">NB</td><td rowspan="2">–</td><td>>100</td><td>NB</td><td>0.9 +/- 0.2</td><td>–</td></tr>
<tr><td>Linker 4 (PI3P)</td><td>–</td><td>–</td><td>NB</td><td>–</td></tr>
<tr><td>Linker 4 R649* (PI5P)</td><td>–</td><td>–</td><td>–</td><td>–</td><td>–</td><td>–</td><td>NB</td><td>–</td></tr>
<tr><td>Linker 2 R296* (PI5P)</td><td>–</td><td>–</td><td>–</td><td>–</td><td>–</td><td>–</td><td>–</td><td>NB</td></tr>
</table>

**Extended Data Table 2: Dissociation constants ($K_D$ in µM) of interaction of wild type and mutant recombinant hUHRF1 proteins with different lipid ligands as determined by microscale thermophoresis.**

UBL-TTD-PHD-SRA: hUHRF1 (aa 1-619); Linker 4: hUHRF1 (605-675); RING: hUHRF1 (675-793); TTD: hUHRF1 (aa 126-285); Linker 2: hUHRF1 (aa 286-306); PHD: hUHRF1 (aa 301-376); hUHRF1 PBR*: K644A, K646A, K648A, R649A, K650A, S651A; hUHRF1 R296*: R296A; hUHRF1 R296R649*: R296A, R649A; hUHRF1 D142*: D142A; hUHRF1 D142R649*: D142A, R649A; hUHRF1 TTD*: Y188A, Y191H; hUHRF1 PHD*: D334, D337A. NB, not binding; nd, not determined.

| | **di-C 16:0 PI5P** | **di-C 16:0 PI3P** | **di-C 16:0 PE** |
|---|---|---|---|
| hUHRF1 (UBL-TTD-PHD-SRA-RING) | 4.7 +/- 0.5 | 5.5 +/- 1 | 3.8 +/- 0.3 |
| UBL-TTD-PHD-SRA | NB | – | – |
| Linker 4 | 2.8 +/- 0.4 | 5.8 +/- 0.2 | NB |
| RING | NB | – | – |
| TTD | NB | – | – |
| Linker 2 | NB | – | – |
| PHD | NB | – | – |
| hUHRF1 PBR* | NB | NB | – |
| hUHRF1 R296* | 4.6 +/- 0.8 | – | – |
| hUHRF1 R296R649* | >80 | 2.3 +/- 0.5 | – |
| hUHRF1 D142* | 59.9 +/- 13 | – | – |
| hUHRF1 D142R649* | NB | 5.8 +/- 2 | – |
| hUHRF1 R649* | 2.6 +/- 0.3 | – | – |
| hUHRF1 TTD* | 4.3 +/- 1.7 | – | – |
| hUHRF1 PHD* | 4.9 +/- 1.3 | – | – |

| | **di-C 16:0 PI5P/DMPC/DHPC bicelles** | **di-C 16:0 PI5P/DHPC bicelles** |
|---|---|---|
| hUHRF1 | NB | 98.7 +/- 25.3 |
